# A three-compartment microfluidic platform for investigating signal transmission in the human sensory pathway^†^

**DOI:** 10.64898/2026.01.06.698004

**Authors:** Blandine F. Clément, Trui Osselaer, Claire Zhang, Giacomo Paccagnan, Tobias Ruff, János Vörös

**Author notes:** These authors contributed equally to this work (Blandine F. Clément, Trui Osselaer, Claire Zhang).

## Abstract

Neuropathic pain remains a significant challenge due to limited understanding of sensory signal transmission mechanisms along the sensory pathway. The sensory pathway involves peripheral nociceptors in the dorsal root ganglia (DRG) that transmit signals from skin to the central nervous system via dorsal horn neurons. Current *in vitro* models lack the compartmentalization and resolution needed to investigate signal modulation at distinct anatomical sites along this pathway. Here, we developed a three-compartment microfluidic platform combining human induced pluripotent stem cell (iPSC)-derived sensory neurons (hiSNs) with human primary epidermal keratinocytes (HPEKs) and iPSC-derived dorsal horn neurons (hiDHNs) in a spatially organized arrangement. The platform integrates polydimethylsiloxane (PDMS) axon-guiding microstructures with high-density microelectrode arrays (HD-MEAs), enabling single-axon electrophysiological recordings and sub-cellular level stimulation. We established viable co-cultures maintained for up to six weeks and characterized spontaneous activity across all conditions. Keratinocytes increased the number of active sensory neuron axons and their firing rates, demonstrating peripheral modulation of neuronal activity. Systematic frequency-dependent electrical stimulation revealed low-pass filtering properties at sensory neuron somata, with filtering characteristics modulated by co-culture with keratinocytes. This platform enables compartment-specific investigation of signal processing in the human sensory pathway and provides a tool for studying neuropathic pain mechanisms and testing potential therapeutics.

## Introduction

Chronic pain is a debilitating and increasingly prevalent condition affecting more than 20% of the population worldwide, with neuropathic pain representing one of the most challenging forms to treat effectively ^1^. The inadequacy of existing treatments highlights the urgent need for novel therapeutic strategies, which require a deeper understanding of the complex pathophysiological mechanisms underlying neuropathic pain.

Pain processing is based on a coordinated communication between the peripheral and central nervous systems through a well-defined anatomical pathway. Sensory neurons are pseudounipolar cells with their soma located in the dorsal root ganglia (DRG), exhibiting a dual axonal projection through a T-shaped bifurcation: one branch extends toward the periphery to innervate skin and other tissues, while the other projects centrally to the spinal cord dorsal horn (DH) to relay neuronal excitability to ascending brain pathways through synaptic transmission at the first sensory synapse ^2,3^. Neuropathic pain pathophysiology involves multiple levels of dysfunction: peripheral sensitization at nerve terminals, altered signal processing at the T-junction, and central sensitization in spinal cord circuits ^4^. The complexity of these interactions across different anatomical compartments requires advanced experimental approaches to dissect individual contributions to pain signaling.

The traditional view of nociception as a purely neuronal process has evolved with growing evidence that non-neuronal cells actively participate in pain signaling. In the skin, epidermal keratinocytes function as environment-sensing cells that can mediate nociceptor sensitization and encode noxious stimuli through various sensory receptors that, after stimulation by different modalities (thermal, mechanical, chemical) ^5–10^, release signaling molecules, activating nearby nerve endings. The interaction between keratinocytes and sensory neurons can be further modulated by inflammatory mediators as in the case of small fiber neuropathy ^11^ or by an increased production of nerve growth factor (NGF) contributing to peripheral hyperalgesia ^12,13^.

Not only is peripheral target tissue a key site in the sensory pathway, but the first synapse between DRG and DH neurons also represents a critical processing point. Indeed, at the central level, nociceptive signals undergo modulation through complex excitatory and inhibitory mechanisms ^3,14^. This synaptic interface is essential for understanding pain processing because it serves as the gateway for sensory information entering the central nervous system. Dysfunction at this level contributes significantly to neuropathic pain development through altered excitation-inhibition balance, reduced GABAergic tone, and pathological synaptic plasticity ^15,16^ which makes the DRG-DH connectivity a crucial mechanism of central sensitization, responsible for some pain states. The complexity of dorsal horn circuitry requires sophisticated experimental approaches to understand how these circuits process sensory information and become dysfunctional in chronic pain.

Additionally, emerging evidence suggests that DRG somata serve important analgesic functions beyond their traditional role as metabolic support centers ^17,18^. Indeed, the clinical relevance of DRG modulation was furthermore demonstrated by the success of dorsal root ganglion stimulation for treating chronic neuropathic pain ^19,20^. Their peculiar pseudounipolar arrangement potentially creates an inhibitory somatic membrane potential, making this component a low-pass filter for action potential propagation, with filtering properties that differ between nociceptor subtypes ^21,22^. This local inhibitory system could be targeted pharmacologically to modulate pain signaling at the ganglia level, representing a potential therapeutic approach that bypasses central nervous system side effects. However, studying DRG somata contributions to pain processing remains technically challenging due to difficulties in selectively accessing the somatic compartment while preserving axonal function and maintaining physiological cellular interactions.

Finally, it is important to study skin-nerve interactions when aiming for a better understanding of pain mechanisms and developing targeted therapies. Yet the functional interactions between nociceptive nerve endings and skin cells remain challenging to study due to the small diameter of nerve endings and difficult experimental accessibility ^23^.

While current skin-on-chip platforms successfully recapitulated epidermal barrier function, stratification, and inflammatory responses ^24,25^, they typically lack neuronal components essential for studying nociceptive signaling. Several research groups have developed microfluidic-based co-culture systems combining keratinocytes with sensory neurons to investigate skin-nerve interactions ^23,26–28^ to enable specific subcellular compartment interventions. Others have incorporated human induced pluripotent stem cell (iPSC)-derived sensory neurons to overcome limitations associated with animal-derived cells ^29,30^. Most recent developments including re-innervated skin models ^31^ and even 3D innervated epidermal-like layers ^32^ have achieved physiologically relevant cellular architecture with enhanced barrier function and anatomical innervation. Yet, they face technical constraints including limited spatial control over axonal growth or inadequate electrophysiological recording capabilities impeding detailed neurophysiology investigations.

Similarly, several models of the DRG-DH connectivity have been developed to study synaptic transmission mechanisms ^33^. Microfluidic co-culture models have successfully recapitulated the first sensory synapse using postnatal mouse sensory neurons, demonstrating salient features of *in vivo* physiology including glutamatergic signaling and calcium responses in DH neurons following DRG stimulation ^34^. Advanced microphysiological systems have demonstrated morphine-sensitive synaptic transmission in bioengineered models of peripheral nerve connected to spinal cord, showing that mechanistically distinct analgesics differentially modulate synaptic function ^35^.

Despite the importance of studying these anatomical compartments in their appropriate cellular environments, current experimental approaches face significant limitations. Recent advances in microfluidic compartmentalization and high-density microelectrode arrays (HD-MEAs) address the limitations of existing platforms by enabling measurement of spontaneous activity and stimulation-induced activity of single axons to better characterize signal modulation in topologically confined small circuits of neurons ^36–39^. The integration of multiple cellular compartments within a single platform enables investigation of complex cellular interactions while maintaining the ability to manipulate and record from individual components.

In this study, we developed a three-compartment microfluidic platform that integrates human iPSC-derived sensory neurons (hiSNs), human primary epidermal keratinocytes (HPEKs) and human iPSC-derived dorsal horn neurons (hiDHNs). This approach addresses the critical need for human-relevant models by incorporating human iPSC-derived neurons alongside human primary skin cells. By employing polydimethylsiloxane (PDMS) microstructures for controlled axon-guidance in combination with HD-MEA, we leverage the high spatiotemporal resolution readout capability offered by the technology to obtain a near-complete coverage of confined circuits of neurons, enabling the study of how specific intervention (electrical stimulation, target cell type addition) modulate network activity ^40^. Our approach could enable comprehensive investigation of peripheral skin-neuron interactions, central synaptic transmission, and potential analgesic mechanisms at the DRG soma level within a single integrated platform. By incorporating all three main surrogates of the sensory pathway components, our platform aims at investigating how dysfunction in one compartment affects signaling throughout the entire pathway. The ability to selectively stimulate and record from different compartments enables the dissection of the individual contributions of each cell type to both normal sensory processing and pathological pain states.

## Results and discussion

### A compartmentalized microfluidic platform integrating human sensory pathway core components

We designed and fabricated a platform featuring three distinct culture chambers incorporating core components of the sensory pathway (“SKIN-DRG-DORSAL HORN”) (Fig. 1A) via the co-culture of hiSNs with two additional cell types, HPEKs and hiDHNs, in the following linear arrangement: HPEKs-hiSNs-hiDHNs. The microfluidic device was designed to enable interaction of hiSNs with HPEKs on one side and hiSNs with hiDHNs on the other side while maintaining individual cell type mid-term viability in a co-culture system. As distinct cell types require different optimal culture conditions (culture medium, substrate coating), separate culture chambers are necessary. The device comprises a polydimethylsiloxane (PDMS)-based bottom layer axon-guiding microstructure, on which two upper PDMS compartmental wells are placed to facilitate viable co-culture via three distinct culture compartments (HPEKs, hiSNs, hiDHNs) (Fig. 1B-C(i)-D(i)). Central to this design, the “DRG” compartment is modeled by hiSNs growing axons inside the axon-guiding microstructure featuring five identical parallel circuits, projecting axon growth towards each side as in the human sensory pathway where primary sensory neurons project distal axons to peripheral tissues on one side and proximal axons to the dorsal horn on the other side. Additionally, as sensory neurons convey information through electrical signaling, this *in vitro* platform is integrated with a high-density microelectrode array (HD-MEA) to enable recording of both spontaneous and stimulation-based electrophysiological activity from hiSNs with single axon resolution. Each circuit features a central seeding well hosting a hiSN spheroid (Fig. 1E). From each side of this central seeding well, 15 axon-guiding microchannels extend outward, beginning with a narrow constricting region approximately 3 *µ*m-wide. Their confined geometry enhances the signal-to-noise ratio for electrophysiological recordings and improves axon-level stimulation capabilities. On one side, microchannels extend into a well (“SKIN” compartment) containing HPEKs in the same plane (in-plane). This compartment includes anti-backgrowth features to minimize axons returning into the microchannels. On the other side, microchannels terminate in specialized microwells (Fig. 1E, right panel), that direct axonal growth upward at a 90-degree angle to an “off-plane” compartment (“D-HORN”) hosting hiDHN spheroids. This vertical growth design prevents hiDHN axons from entering the microchannels and interfering with measurements of hiSN action potential. Diffusion microwells positioned along the axon-guiding microchannels facilitate the diffusion of nutrients, oxygen, and other compounds such as dyes, except where the upper wells are attached. Each microchannel is spanned by a high density of electrodes, enabling measurement of individual action potentials from axons contained within the microchannels with precise spatial and temporal resolution. This allows for detailed characterization of signal propagation. The complete design specifications can be found in Fig. S1 (Supporting Information). Designs for the spatial distribution of microchannels in each circuit were not changed from previous designs, and assure that when laid out on top of the sensing area, across their width, each microchannel cannot be measured by more than one electrode, and each electrode cannot measure more than one microchannel. Overall, this platform enables comprehensive analysis of electrophysiological signals in a model of the sensory pathway, allowing high-resolution and high-throughput data collection through the integration of five parallel axon-guiding circuits on a single HD-MEA.

**Fig. 1.**
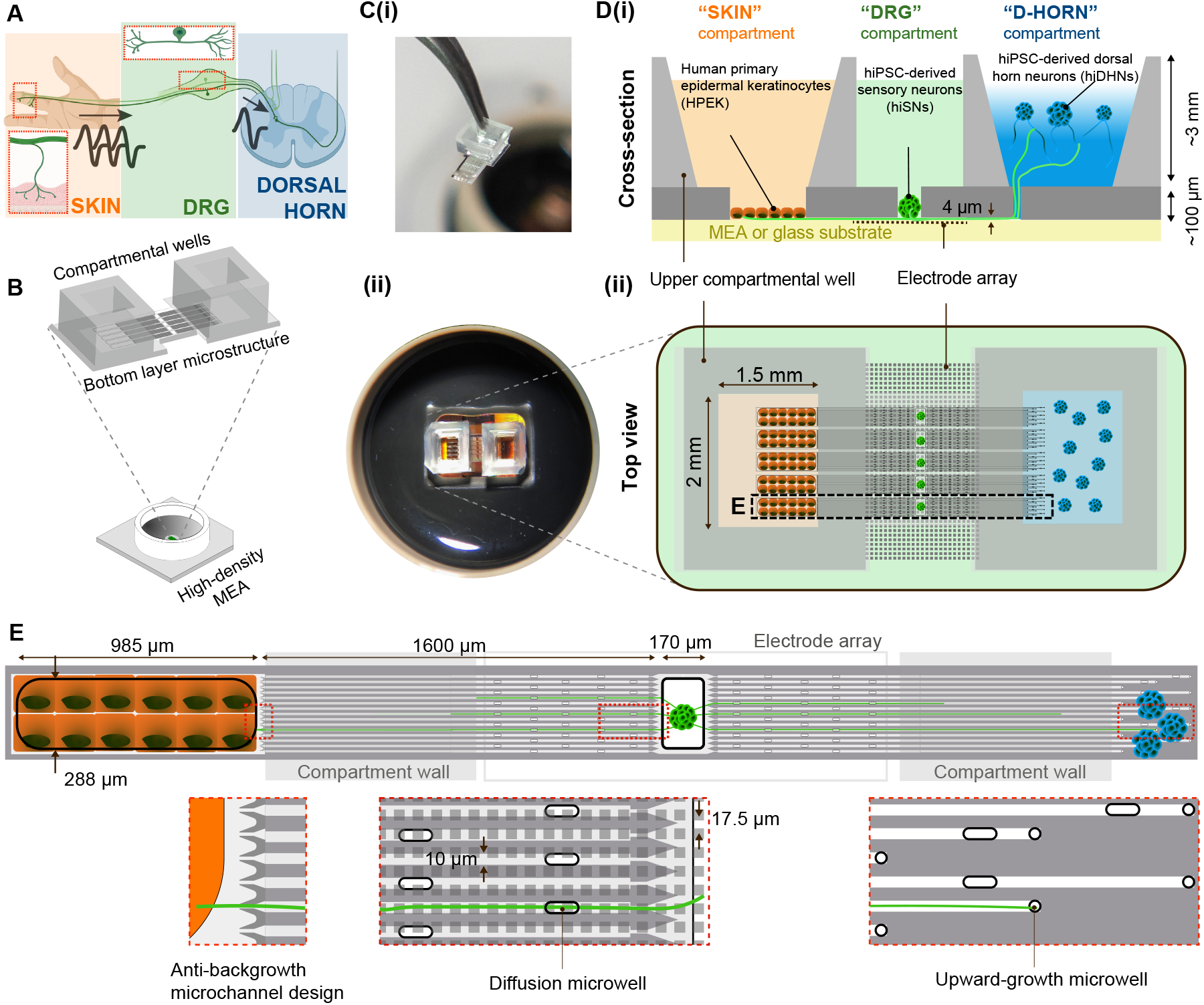
Overview of the compartmentalized microfluidic model of the sensory pathway. **A**: Simplified illustration (generated with BioRender) of the signal propagation in the anatomic sensory pathway, highlighting three core components: peripheral tissue (orange; SKIN), primary sensory neurons (green; DRG), and dorsal horn of the spinal cord (blue; DORSAL HORN). **B**: The platform consists of a microfluidic device placed onto a high-density microelectrode array (HD-MEA). The microfluidic device contains a bottom layer PDMS microstructure featuring five identical parallel axon-guiding circuits on top of which two upper PDMS compartmental wells are placed on each side of the circuits, creating three distinct cell culture chambers. **C**(i): Photography of the microfluidic device with only one upper well mounted on one end. (ii): Photography of the platform through a stereo-microscope. **D**: Cross-section (i) and top (ii) view of the three-compartment model. The two upper peripheral wells allow compartmentalization of the co-cultured HPEKs “in-plane” on one side (“SKIN” compartment) and hiDHNs “off-plane” on the other side (“D-HORN” compartment) of the hiSNs seeded in the center (“DRG” compartment) of the circuits. **E**: Schematic of one axon-guiding circuit with dimensions. Left subpanel shows the design where the axon-guiding microchannels enter the “SKIN” compartment intended to prevent backgrowth from axons. Middle subpanel shows diffusion microwells along the microchannels to facilitate nutrient exchange. Right subpanel shows microwells at the end of the microchannels at the “D-HORN” side enabling growth of the hiSN axons in the upper chamber.

### Optimization of keratinocyte seeding and viability in the “SKIN” compartment

After building the compartmentalized platform, we investigated the optimal culture conditions of HPEK within the constraints posed by the small dimensions of the upper wells. Viability is needed to promote interactions with hiSNs for a sufficiently long working window to allow experimental measurements (Fig. 2). We first tested three variables for HPEK culture viability: initial seeding density, passage number, and substrate material. HPEK culture viability was assessed by surface coverage based on fluorescently labeled live cells. Overall, surface coverage consistently dropped between day *in vitro* (DIV) 7 and DIV 14 for all conditions. Seeding at the conventional initial seeding density of 4,000 cells/cm^2^ resulted in highly variable health of HPEK cultures at DIV 7, while seeding at higher density (40,000 and 80,000 in the well compartment ensured more reliable surface coverage at DIV 7 that would drop systematically at DIV 14. Seeding at lower density allowed for maintaining viability and greater than 50% surface coverage. It is worth mentioning that diluting the cell suspension to a target concentration in a reproducible way for the small volume of the compartment poses challenges, and could be the main factor of variability. Finally, we opted for an initial seeding concentration of 4,000 cells/cm^2^ for our co-culture protocol. Investigating the surface coverage of HPEKs seeded at different passage numbers showed a significant decrease between passage 5 (P5) and P7 at DIV 7; we confirmed that seeding consistently at an early passage number of P5 is not only beneficial for reproducibility reasons, but it also indicated a higher proliferative ability (Fig. S2B, Supporting Information). Finally, we saw that there was no significant difference between seeding on a Matrigel-coated PDMS substrate compared to uncoated PDMS or glass (Fig. S2A, Supporting Information). Testing Matrigel-coated PDMS as substrate mainly aimed at optimizing the HPEK culture in an “off-plane” interface scenario initially thought for the “SKIN” compartment (Fig. S3, Supporting Information) but was not kept throughout our co-culture protocol for the in-plane strategy due to the absence of any visible improvement, and a longer time needed for axons to grow inside a longer distance to the off-plane surface. Additionally, the in-plane strategy for HPEK co-culture was selected due to a drastic lack of HPEK viability in a Matrigel hydrogel (that would have been required for proper axonal outgrowth and outreach to HPEKs), as opposed to a Matrigel surface coating (data not shown). This is most likely due to the fact that HPEKs, if cultured alone, require a stiff substrate (PDMS, glass), and such stiffness range is preserved when Matrigel is only used as a surface coating. As an additional interesting observation, we saw that the in-plane design overcomes one challenge faced with culturing HPEK inside the space of the compartment, thanks to the inter-circuit walls: viable cells are limited to the border of the culture area, whereas the health of cells in the middle is often compromised first (Fig. S2C, Supporting Information). However, occasionally and locally HPEKs’ confluency even reached a monolayer state (Fig. S2D, Supporting Information).

**Fig. 2.**
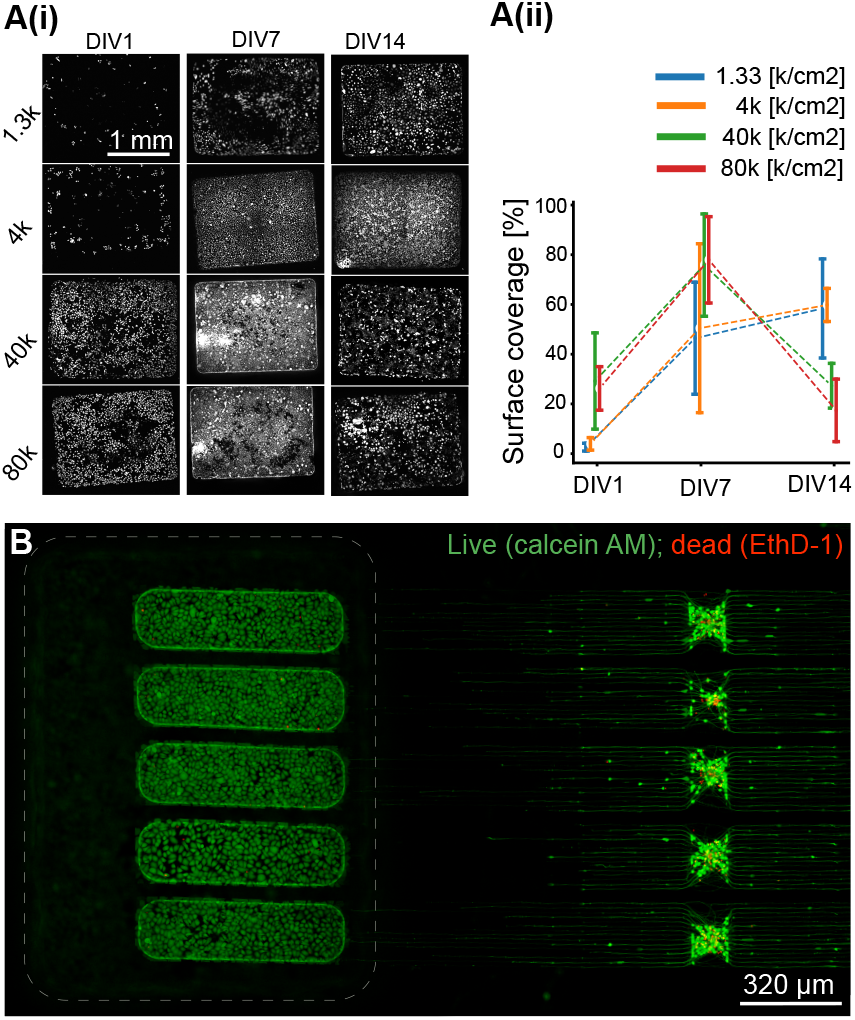
Optimization of HPEK seeding conditions to enable hiSN-HPEK co-culture. **A**(i): Representative fluorescence images for each initial seeding density condition (live cells are stained with Calcein AM). (ii): HPEK surface coverage across days *in vitro* (DIV) for different initial seeding densities. **B**: Live-dead staining of hiSN and HPEK shows viability of both cell types in respective compartments.

### Sequential seeding strategy enables keratinocyte-neuron interactions

Another aspect of optimizing the HPEK culture was to enable a healthy culture interfaced by neurons. Prior attempts of hiSN-HPEK co-cultures were carried out by seeding the two cell types simultaneously (same day, with only a few hours differences). Although no significant difference was found between the growth rate of axons in the presence and absence of HPEKs, it is interesting to observe that all curves corresponding to the HPEK case are above the control curves indicating some potential influence through paracrine signaling (Fig. S2E, Supporting Information). Beside this effect, at the time axons had reached the “SKIN” compartment, HPEKs had already proliferated and crowded the area of the compartment (Fig. 3A(i)). Although they had not reached full confluency yet, it seemed their presence at the interface and most particularly their migration below the lower roof interface with microchannels was enough to form a physical barrier preventing the axons to grown inside the compartment and in some occurrences making the axons grow back into a neighboring microchannel, despite the anti-backgrowth design feature at the channel exit (Fig. 3A(ii-iii). We saw that seeding HPEKs three days later than hiSNs helps axons to grow past the interface between the microchannels and the open compartment (Fig. 3B(i)). Although the fluorescent tags were non discriminative between cell types in some cases, the difference in morphology between axons and HPEKs was enough to assess their interactions. We saw with time-lapsed images that axons transiently “sense” and interact with HPEKs through their growth cone filopodia when the two cells are close-by, but axons eventually find a growing path bypassing HPEKs, and usually end up growing along the lateral compartment walls where the narrow confining roof edge limits the HPEKs’ presence (Fig. 3B(ii-iv) and Fig. S4, Supporting Information). An additional example of sequential seeding shows axons grown around HPEKs at a later stage of HPEK confluency (Fig. S5A(i-iii), Supporting Information). Seeding HPEKs 14 days after (Fig. 3C) was identified as the right protocol as a trade-off between allowing a much higher number of axons grown inside the target compartment as it is the case in the total absence of HPEKs (Fig. S5B, Supporting Information), and starting a co-culture with HPEK early enough to measure how HPEKs affect neuronal activity. After two weeks in co-culture, hiSN axons interfaced nicely with HPEKs, although respective arrangement and morphology of both cell types was variable across circuits and samples.

**Fig. 3.**
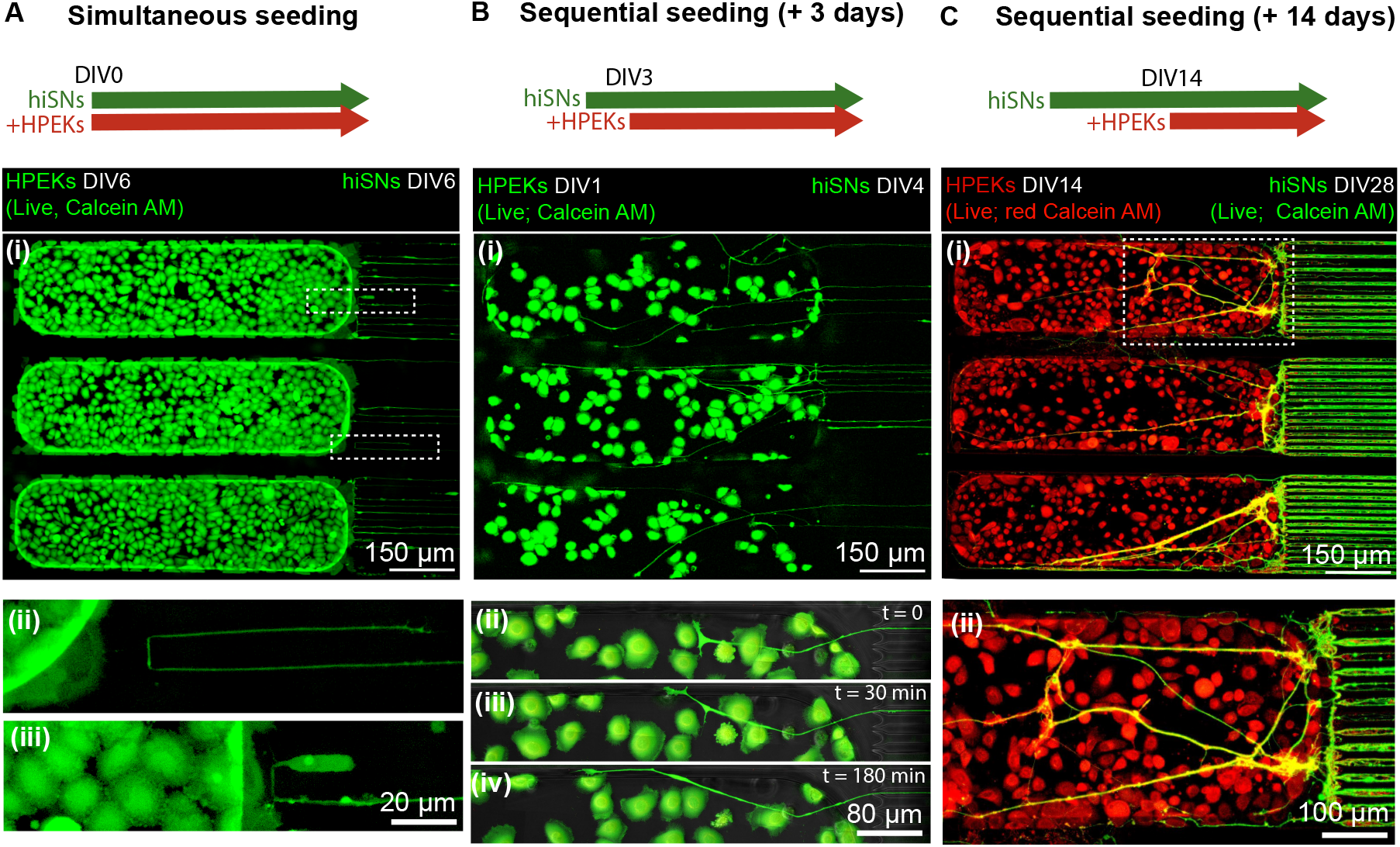
Optimization of HPEK-hiSN co-culture conditions **A-C**(i): Fluorescence images of the “SKIN” compartment with or without incoming axons from microchannels when HPEKs were seeded in the same time as hiSNs (A); three days after hiSN seeding (B); 14 days after hiSN seeding (C). **A**(ii-iii): Zoomed in views of A(i) showing axons growing back due to physical barrier formed by HPEKs. **B**(ii-iv): Zoomed in time-lapsed views of B(i) showing an axon invading the “SKIN” compartment at time = 0, 30 and 180 minutes. **C**(ii): Zoomed in view of C(i) showing axons grown between and on top of HPEKs.

### Integration of dorsal horn neurons completes the three-compartment sensory pathway model

After optimizing the compartmentalized co-culture of HPEKs and hiSNs, adding hiDHNs established a viable co-culture of the three cell types resembling the organization of anatomical components of the sensory pathway, in the form of a three-chamber co-culture (Fig. 4A(i)). hiSN spheroids were seeded in the central seeding wells between the two upper wells of the platform (Fig. 4A(iii). As evident in Fig. 4A(ii), the hiSNs interface with the HPEKs. In order to extend the model towards the first synaptic target in the central nervous system, we integrated hiDHNs into the co-culture system. When co-culturing hiDHNs, we need to consider unambiguous electrophysiological interaction with the system. To this purpose, it is important that only hiSN axons grow in the axon-guiding microchannels, while hiDHN axons remain in the “D-HORN” compartment. Therefore, an off-plane design was elected at the “D-HORN” side, rather then an in-plane compartment as at the “SKIN” side. The microwells at the end of the microchannels in the off-plane design allowed upward growth of hiSNs (Fig. 4B(ii) in green) given that hiSN axons would grow until the microchannel ends (Fig. 4A(iv), while simultaneously reducing the likelihood of hiDHNs growing downward. However, when hiDHN spheroids were seeded directly on top of the microstructures, axons tended to seek out the irregularities in the surface at the microwells, subsequently growing into the microchannels (Fig. S6, Supporting Information). To mitigate this, hiDHN spheroids were seeded in Matrigel, providing a three-dimensional (3D) growth environment that minimized the impact of surface irregularities and fostered a more physiological environment. Lastly, hiDHNs were seeded two weeks after hiSNs, to allow hiSN axons to fill the microchannels, thereby hindering entry of hiDHN axons into the microchannels. The described procedure enabled hiSNs and hiDHNs to interface in the 3D “D-HORN” compartment 7 to 14 days after hiDHN co-seeding (Fig. 4B). Potentially, the volume of Matrigel could be decreased, to allow interaction between neuron types to occur at an earlier time point. This viable co-culture of hiSNs, HPEKs and hiDHNs on HD-MEAs enabled investigation of signal transmission and modulation in the sensory pathway.

**Fig. 4.**
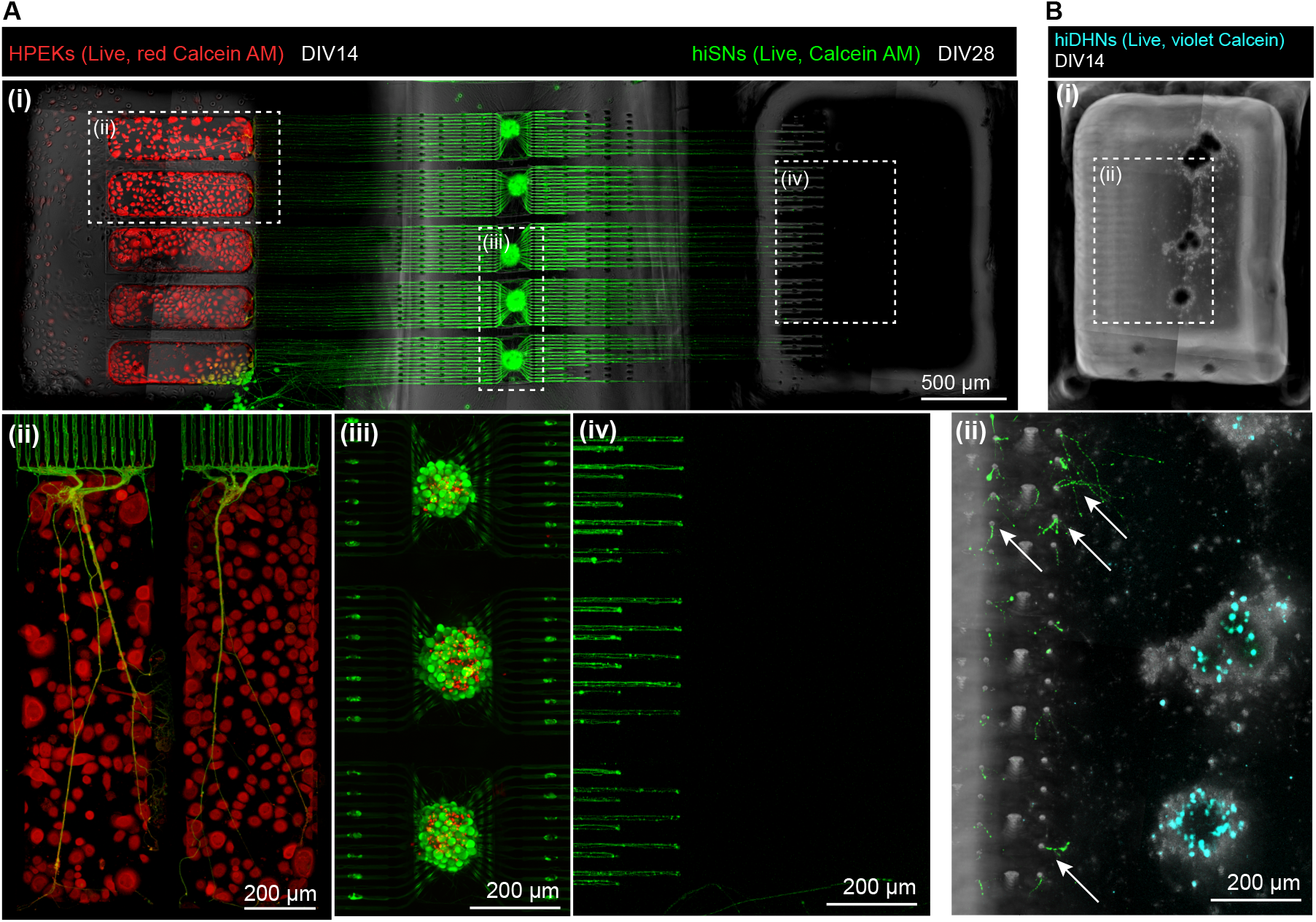
Overview of the compartmentalized co-culture with hiSNs, HPEKs and hiDHNs. **A:** Composite image of the three-compartment co-culture of hiSNs DIV 28 (Live, green Calcein AM) and HPEKs at DIV 14 (Live, red Calcein). Image shows the microchannel plane. (ii) Zoomed-in view of A(i) of the “SKIN” compartment where hiSN axons grew in (green). (iii) Zoomed-in view of (i) on the hiSN somata in the central seeding wells (Live/dead – green/red staining). (iv) Zoomed-in view of the end of microchannels projecting below the “D-HORN” compartment. **B:** Bright-field image of the “D-HORN” compartment shown in A at a different plane showing hiDHN spheroids seeded higher up (off-plane) in Matrigel. (ii) Zoomed-in view of B(i) and higher plane (approx. 150 **µ**m above of A(iv)). The image is a z-projection of all planes from the top of PDMS microstructures where hiSN axons grew upward out of the microwells (white arrows) in Matrigel in 3D (green) and hiDHN spheroids are in blue (violet Calcein).

### Keratinocytes modulate sensory neuron firing rate without affecting spontaneous signal directionality

As outlined in the introduction, this study aimed to characterize the spontaneous activity in an established *in vitro* model of the sensory pathway, and investigate the modulation of this signal due to co-culture with HPEKs and hiPSC-derived DH neurons. To achieve this goal, our platform design enabled distinct recording of spontaneous electrophysiological activity from hiSN axons in each of the 15 microchannels on both sides of the platform. An analysis pipeline was developed to identify single axons in microchannels and further characterize their spontaneous activity over time, with and without co-culture of the two target cell types: HPEKs and hiDHNs (Fig. 5). In general, the results demonstrate that our model successfully maintains viable neural activity for up to 6 weeks in culture and reveals significant modulatory effects of HPEKs on spontaneous activity patterns. As illustrated in Fig. 5D and E, maximum spontaneous activity was observed in all conditions at DIV 28, both in terms of the number of active axons and the firing rate of these axons. Furthermore, the measurements clearly demonstrate the viability of the system up to DIV 42, 4 weeks after co-seeding of HPEKs and hiDHNs in the respective conditions. Moreover, the effect of co-culture is evident in the increased number of active axons, with the most pronounced effect resulting from the addition of HPEKs. Indeed, the firing rate of the hiSNs in co-culture with HPEKs was notably increased. This finding is consistent with recent reports regarding the role of keratinocytes in nociception, and their potential contribution towards peripheral sensitization ^6^. The slight observable decrease in number of active axons and firing rate in all conditions from DIV 14 to DIV 21 could be attributed to the switch from Senso-MMx2 Maturation Media to a custom-made medium after DIV 14, as described in the experimental section. Fig. 5D further shows that co-culture with HPEKs introduced asymmetry into the system, with a higher number of active axons identified at the “SKIN” side (indicated by the larger values for the darker color symbols). As demonstrated in Fig. 5G, signal propagation in all conditions is predominantly from the center to periphery, indicating that propagation along the physiologically expected direction, *i*.*e*., from the HPEKs to the hiSN somata, is not induced. This directionality pattern could be attributed to an incomplete differentiation of the hiSNs, causing them to lack the pseudounipolar morphology or ion channel distribution essential for peripheral-to-central signal directionality ^2,41^. Interestingly, at DIV 7, propagation on both sides of the platform is more equally distributed in both directions. The apparent more peripheral origin of spikes at this early stage could be attributed to responses to mechanical cues present in the microchannels during axonal growth ^42^. Furthermore, at DIV 21, we observed an increase in signals originating from the “D-HORN” side of the model (light blue symbol in Fig. 5G), possibly due to signaling between the hiDHNs and hiSNs. However, since this effect did not persist, it may not be functionally meaningful. As evident from Fig. 5F, the conduction speed of the hiSNs decreased over time, which could indicate differentiation towards a subtype with lower conduction speed. Co-culture with HPEKs or hiDHNs did not appear to affect the conduction speed of the hiSNs. Additionally, firing rate and conduction speed were not correlated, suggesting that spontaneous activity operated independently from the structural properties that determined conduction speed (Fig. S8, Supporting Information). The more detailed distributions of firing rate and conduction speed values across DIV are presented in Fig. S7 (Supporting Information). Despite the promising results, a limitation of our current model is the lack of directional signal propagation that would more accurately reflect *in vivo* conditions. Future work should explore co-culture of hiSNs with satellite glia, which has been reported to advance the maturation of DRG neurons, including the development of a pseudounipolar morphology ^41^. Additionally, introducing peripheral stimuli to the system using sensitizing compounds such as capsaicin could potentially induce more physiological relevant signal propagation patterns ^23,27^. Regardless, our platform successfully enables the characterization of spontaneous activity in this sensory pathway model, allowing for the identification of modulatory effects of both hiDHNs and HPEKs on neural signaling. These findings suggest that keratinocytes play an active role in modulating sensory neuron activity, which could have important implications for understanding peripheral sensitization mechanisms and inflammatory pain conditions.

**Fig. 5.**
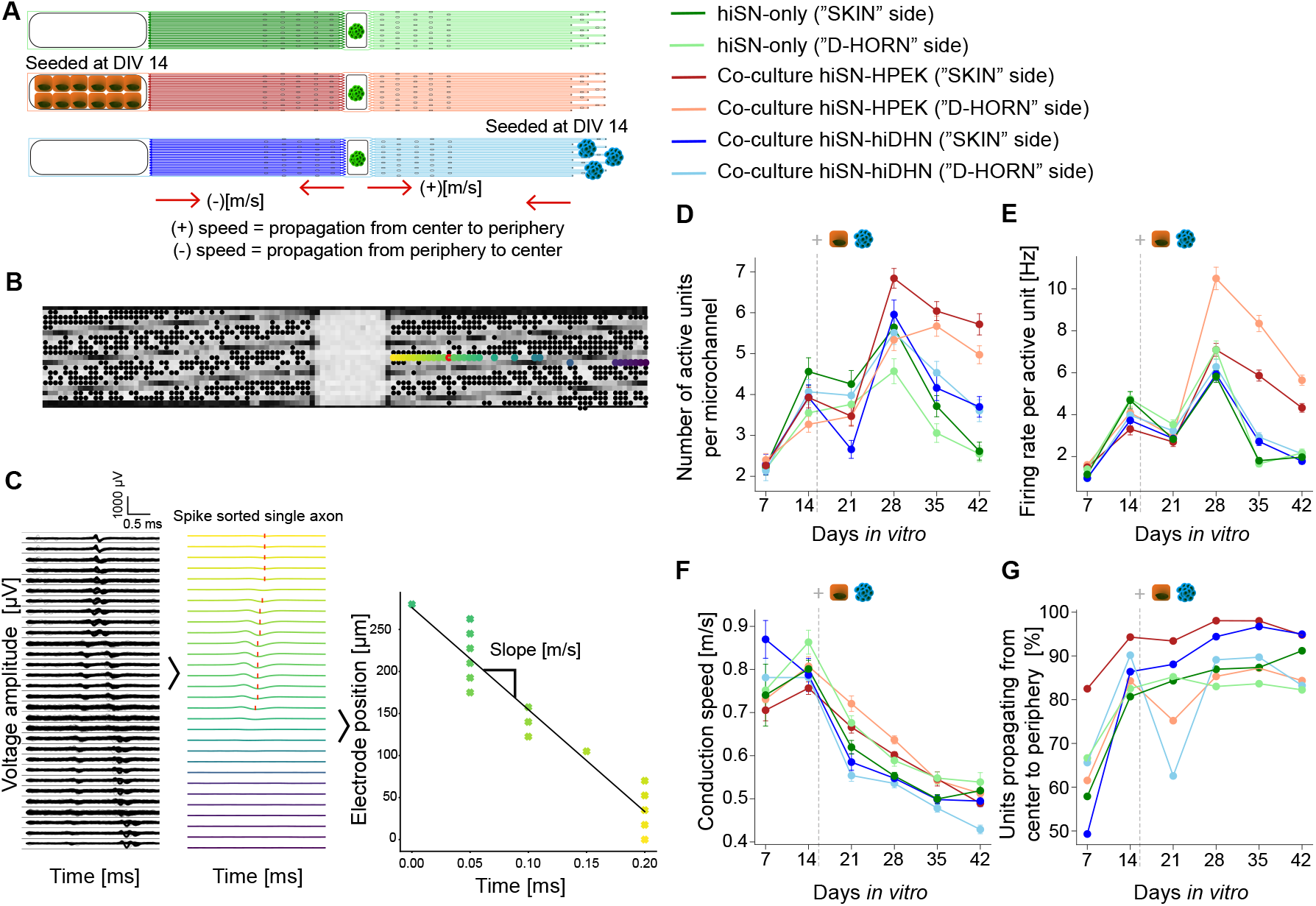
Characterization of spontaneous activity in the model. **A:** Overview of the co-culture experimental conditions. **B:** Spatial distribution of the electrodes used to measure the propagation of a spike within a microchannel in C. **C:** Representative example of measured propagating spikes in both directions along pre-identified microchannels before and after spike sorting. Waveforms are shown for one sorted axon, based on the mean waveforms along the recording electrodes in the microchannel. **D:** Number of active axons per microchannel, means and standard error of means over microchannels are plotted. **E:** Firing rate per active axon, means and standard error of means over active axons are plotted. **F:** Conduction speed, means and standard error of means over active axons are plotted. **G:** Percentage of active axons propagating from center to periphery. (Green: hiSN-only, orange: hiSN-HPEK co-culture, blue: hiSN-hiDHN co-culture; darker colors indicate axons leading to “SKIN” compartment, lighter colors are used for axons on the side of “D-HORN” compartment.)

### Frequency-dependent filtering properties of sensory neuron somata revealed by systematic electrical stimulation

As discussed in the introduction, sensory neurons have been suggested to function as low-pass filters, with the filtering occurring specifically at the T-junction of the somata ^21,22^. To investigate these filtering properties in our model, we developed a comprehensive stimulation protocol that allowed us to stimulate hiSN axons at the “SKIN” side with different pulse frequencies, while analyzing the stimulation-induced responses along the pathway. Our experimental design enabled us to distinctly isolate the effects occurring in the somatic region from potential effects occurring in the axons. In general, our results provide compelling evidence for the low-pass filtering function of sensory neuron somata, with this effect being modulated by co-culture conditions. As illustrated in Fig. 6, we observed clear frequency-dependent effects on signal transmission across the pathway, from axons to somata. To isolate the role of the somata specifically, we stimulated each microchannel separately, and identified stimulation-induced responses at two key locations: first at one control electrode in the stimulated microchannel close to the central compartment (before the somata), and second at a target electrode in each of the 15 microchannels at the “D-HORN” side (after the somata) (Fig. 6A and Fig. S9 (Supporting Information). This approach allowed us to directly compare signal propagation before and after the somatic region. The low-pass filtering properties of the somatic region are clearly evident in Fig. 6C, where the number of activated microchannels measured at the target electrodes decreased significantly with increasing stimulation frequency, while remaining approximately stable at the control electrodes detecting responses before the somatic region. This indicates that the transmission of higher-frequency pulse trains is selectively blocked in the somatic region of the hiSNs. Notably, this filtering effect was more pronounced in the co-culture conditions, particularly with HPEKs, suggesting that the addition of keratinocytes potentially induces some differentiation in the hiSNs that enhances their filtering properties. Further evidence of the low-pass filtering effect is demonstrated in Fig. 6E, which shows the number of stimulation-induced responses per activated microchannel following the same frequency-dependent pattern as the activated microchannels. However, it should be noted that when interpreting this metric, a stimulation-induced response representing a single axon might be detected twice, as responses are aggregated over separate stimulations, and stimulation of neighboring microchannels is possible. As shown in Fig. 6D, the normalized response probability, defined as the percentage of transmitted pulses in a pulse train, decreased significantly for higher stimulation frequencies at the target electrodes for all conditions. This frequency-dependent filtering effect has previously been observed both *in vitro* and *in silico*, where the T-junction was identified as the location of filtering ^21,43,44^. Our observations suggest that by DIV 28, a substantial proportion of hiSNs have differentiated towards a functional phenotype capable of exhibiting such filtering properties. The filtering function of hiSN somata is further confirmed by analyzing the point of conduction failure, the last stimulation pulse (in a train of pulses) to elicit a response, as shown in Fig. 6F. Conduction failure occurred significantly earlier with increased frequency for all conditions at the target electrodes, while remaining stable at the control electrodes. Our strategical use of short pulse trains of 100 pulses ensured that conduction failure in the axon was limited, thus maximizing the observation of an effect due to the somatic region. Indeed, we have shown in a previous study that conduction failure in the hiSN axons occurred on average after 100 stimulating pulses for our frequencies of interest ^39^. Finally, similar frequency-dependent effects on the stimulation-induced responses were found for stimulation at the “D-HORN” side, indicating symmetry in the filtering properties of the hiSNs (Fig. S9, Supporting Information). These findings are consistent with recent work by Hao et al. ^22^, who demonstrated that DRG neurons control nociceptive input to the central nervous system through a GABAergic system that contributes to low-pass filtering. Importantly, they found this filtering mechanism to be more efficient in C-fibers compared to A-fibers, which aligns with earlier patch-clamp experiments showing failure in pulse transmission at lower frequencies for C-fibers ^21^. The differential filtering effect observed in our co-culture conditions, particularly with HPEKs, suggests that peripheral targets may influence the development of specific hiSNs subtypes or modulate the communication between somata themselves. Despite the robustness of our findings, certain technical limitations must be acknowledged. These include the potential stimulation of neighboring microchannels as mentioned previously, and the sensitivity of response detection to electrode selection. While stimulation, target, and control electrodes are selected based on an activity map as described in the experimental section, this approach does not ensure capturing of all responses. One could address this by selecting multiple control or target electrodes per microchannel, though this would significantly increase computational demands. Additionally, we were unable to test frequencies exceeding 100 Hz due to hardware constraints of the platform, which limits our ability to fully characterize the upper boundaries of this filtering effect. In conclusion, our results provide strong evidence for the filtering role of neuron somata in sensory signal transmission, with this function being modulated by co-culture conditions. The pronounced filtering effect observed in the HPEK co-culture suggests that peripheral targets like keratinocytes may play an important role in shaping the signal processing capabilities of sensory neurons. These findings have significant implications for understanding sensory processing in both normal and pathological conditions, potentially offering new insights into the development of therapies for neuropathic pain and related disorders. Future research should focus on identifying the specific mechanisms by which sensory neurons communicate among themselves to coordinate this filtering function, as well as analyzing how different sensory neuron subtypes contribute to the overall filtering properties of the sensory pathway.

**Fig. 6.**
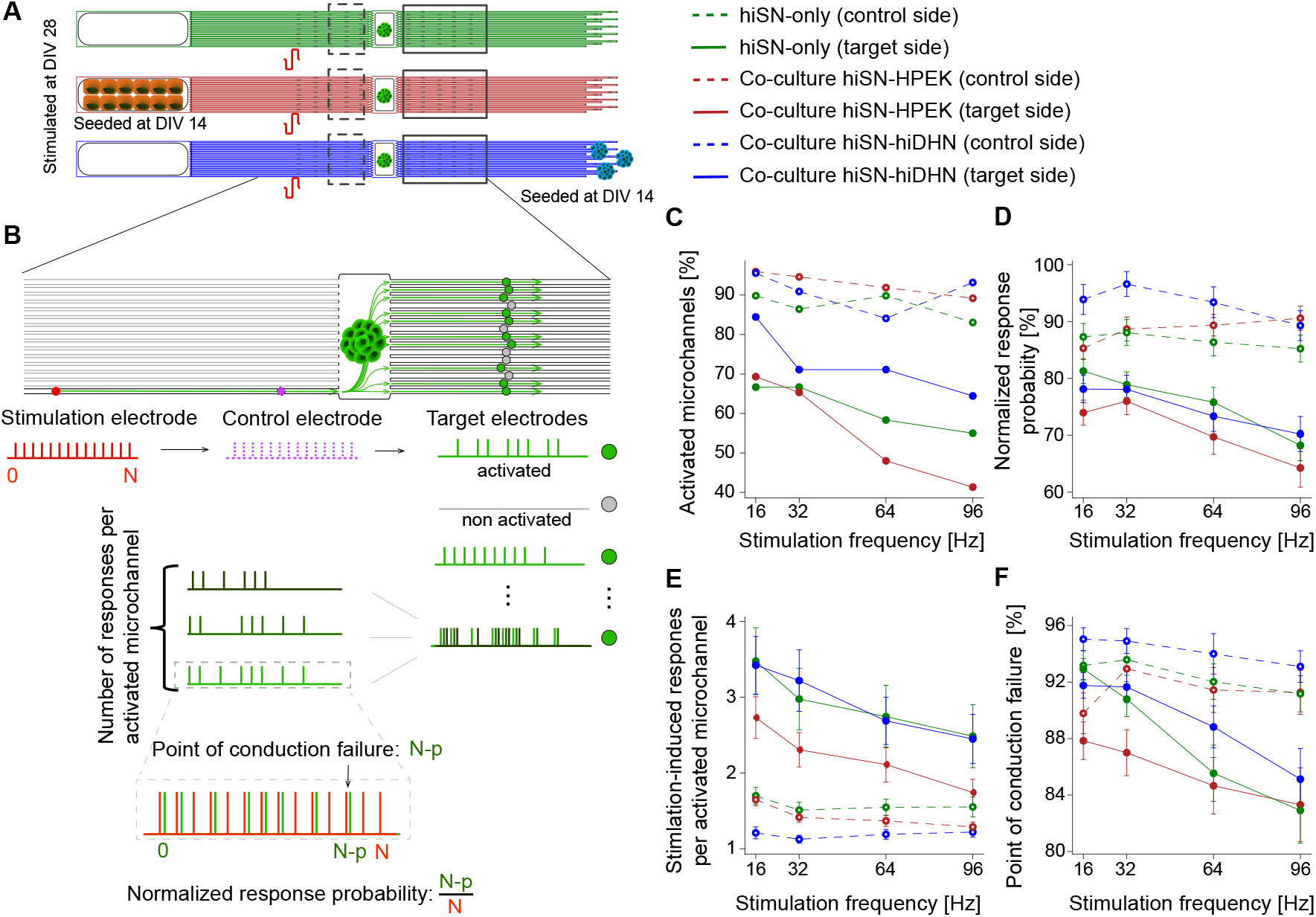
Stimulation-induced responses across hiSN somata are frequency-dependent. **A:** Overview of co-culture conditions stimulated at different frequencies. **B:** Schematic of extracted metrics to quantify and characterize stimulation-induced response transmission. **C:** Percentage of microchannels that are activated, *i*.*e*., have at least one detected response. **D:** Normalized response probability as percentage of total applied stimulation pulses. **E:** Number of detected stimulation-induced responses per activated microchannel. **F:** Point of conduction failure as percentage of total applied stimulation pulses. All curves show means and standard error of means. There are no error bars for the activtaed microchannels (C) as this is a percent of total X measurements. (Green: hiSN-only, orange: hiSN-HPEK co-culture, blue: hiSN-hiDHN co-culture)

## Conclusions

We developed a three-compartment *in vitro* model of the sensory pathway on a HD-MEA, enabling us to investigate signal modulation along the pathway. Using a compartmentalized PDMS-based microfluidic platform, we established a viable co-culture of human iPSC-derived sensory neurons, human iPSC-derived dorsal horn neurons, and human primary epidermal keratinocytes. The combination of the PDMS axon-guidance microstructure and the HD-MEA enabled precise electrophysiological characterization with the platform at the axonal level. We recorded spontaneous activity of the sensory neurons, characterizing the system and demonstrating the active role of keratinocytes in sensitization. By stimulating the platform and identifying stimulation-induced responses along the pathway, the signal transmission in the somatic region of the sensory neurons was characterized, demonstrating low-pass filtering properties. In order to further improve clinical translatability, glial cells could be added to the model to promote differentiation of the hiPSC-derived sensory neurons ^41^. Future research could identify filtering profiles of sensory neuron subtypes, and investigate how induced pathologies affect these profiles. Furthermore, the platform could have applications in drug screening through administration of compounds to individual compartments ^39^. Thus, we could evaluate the platform’s potential in improving understanding and treatment of neuropathic pain.

## Experimental section

### Substrate preparation

For electrophysiological recordings of *in vitro* neuron cultures, a CMOS-based HD-MEA (MaxOne, flat surface topology with less than 0.3 *µ*m variation, MaxWell Biosystems AG, Switzerland) was used featuring 26 400 electrodes allowing for the simultaneous recording from up to 1024 electrodes at 20 kHz. For staining experiments and microscopy imaging, 35 mm diameter glass bottom dishes (HBST-3522T, WillCo Wells) were used. Dishes were cleaned with isopropyl alcohol (IPA) for 10 minutes and subsequently rinsed three times using sterile ultrapure water. The chips and dishes were coated with 0.1 mg/mL poly-L-ornithine (Sigma-Aldrich, P4957) for at least 30 minutes at room temperature and subsequently washed twice with sterile ultrapure water and dried with a nitrogen gun.

### Microfluidic platform design and fabrication

Polydimethylsiloxane (PDMS) microstructure layouts were designed using AutoCAD (Autodesk). PDMS microstructure designs were fabricated by WunderliChips GmbH (Zurich, Switzerland). Each microstructure features five identical circuits. Each circuit features one central seeding well, the dorsal root ganglion (DRG) well, from which an array of 15 (per side) 10 *µ*m-wide, 4 *µ*m-high, 1600 *µ*m-long axon-guiding microchannels extends on both sides. On one side, these channels are connected to an in-plane “SKIN” compartment. On the other side, these channels end with a microwell of 10 *µ*m diameter, allowing the sensory neuron axons to grow upwards into an off-plane “D-HORN” compartment. Microchannels are connected to the outside liquid medium environment via diffusion microwells except where the PDMS upper wells are mounted, to prevent uncured PDMS from leaking inside during the mounting process (as described below). The microchannels have 3 *µ*m-wide constrictions where they connect to a well to minimize the number of axons that can grow through. The exact design specifications and dimensions are indicated in Fig. S1 (Supporting Information).

PDMS microstructures were manually cut out of a wafer. Individual PDMS upper wells were manually cut from 4-microwell inserts (micro-Insert 4 Well, ibidi, 80406) and were mounted on top of the microstructure at each side around the designed keratinocyte and DH compartments to allow for distinct cell type culture conditions (culture medium, coating). One PDMS well has inner opening dimensions at the bottom of 1.5 mm × 2 mm, such that it fits lateral compartments from the five circuits. To ensure long-term and robust adhesion of compartment wells, each well bottom face was stamped in uncured PDMS (1:10 weight ratio of crosslinker to base, SYLGARD 184 Silicone Elastomer Kit, Dow Chemical), and subsequently stamped on a flat surface until excess PDMS was removed, before being carefully mounted on the microstructure. The samples were then cured at 80 °C for at least 2 hours. After attachment of the upper wells, the PDMS microstructures were cleaned in IPA for 10 minutes in an ultrasonic bath, rinsed three times using sterile ultrapure water, followed by 10 minutes in ultrapure water in the ultrasonic bath, and finally dried using a nitrogen gun. It is essential that both the microstructure and the substrate are perfectly dry for the placement described in the next paragraph.

### Mounting microfluidic device on MEA chip

Microstructures were then manually mounted onto the 3.85 mm × 2.10 mm large sensing area, making sure they were placed in the center aligned to the axes of the rectangular sensing area. The reference electrode surrounding the sensing area was left uncovered. The MEA chip was then filled with phosphate buffered saline (PBS, Sigma-Aldrich, 10010015) pre-warmed to 37 °C ensuring the microstructure and PDMS wells were fully submerged, and desiccated to remove air pockets from within the microchannels. Microstructure placement and adhesion were then assessed by collecting an impedance map of the electrode surface as described previously in ^37^. Briefly, a sinusoidal electrical signal (16 mV peak-to-peak voltage, 1 kHz frequency) was generated using an on-chip function generator. The sinusoidal signal was applied to the circumferential reference electrode of the MEA and the received signal at the microelectrodes was recorded. A highly attenuated signal indicated the electrodes covered by PDMS. Iteratively, desiccation was repeated if the impedance map showed the presence of air pockets trapped inside the microchannels. Finally, after sufficient adhesion, the PBS was replaced with a laminin coating solution (iMatrix-511 silk, Anatomic, AMS.892 012) diluted 1:50 in DPBS(-/-, Sigma-Aldrich, D8537) overnight at 37 °C. Despite all care, sometimes part of the PDMS microstructure was not properly attached to the substrate, *e*.*g*., due to minuscule dirt, leading to uncontrolled undergrowth of axons.

### hiSN and hiDHN spheroid formation

A human iPSC-derived sensory neuron (RealDRG™ Nociceptors, Anatomic Incorporated, cat. 1020) or dorsal horn neuron (RealDHN™ Dorsal Horn Neurons, Anatomic Incorporated, cat. 5020) cryovial was removed from liquid nitrogen storage and was placed into a 37 °C dry bath. Cells were thawed (*<*5 minutes) in the bath until only a small bit of ice was left in the vial. After taking the cryovial into a laminar flow cabinet, the cells were gently transferred into a sterile centrifuge tube. The cryovial was rinsed with 1 mL of DMEM/F12 (Gibco™, 11320033) pre-warmed to 37 °C transferring its content also to the sterile centrifuge tube. 6 mL of warmed DMEM/F12 was added slowly to the cell suspension in the tube, which was then centrifuged at 300 g for 4 minutes. After the centrifugation, the supernatant was aspirated without disturbing the pellet. The cell pellet was resuspended in 1 mL of warm maturation medium to create a homogeneous cell suspension. For the sensory neurons, Chrono™ Senso-MM Maturation Media (Anatomic Incorporated, cat. 1030) was used, for the dorsal horn neurons, DHN-MM Maturation Media (Anatomic Incorporated, cat. 5030). After performing a cell count, the cell suspension was dispensed in one or several wells of a spheroid-forming wellplate (Sphericalplate 5D^®^, Kugelmeiers^®^, SP5D-24W), prefilled with 1 mL of the respective warm maturation medium, with the volume necessary to create spheroids of approximately 250 - 500 cells per spheroid. In each spheroid-forming plate well, containing 750 microwells, a total number of 333 000 neurons was seeded, resulting in spheroids with an average size of 450 cells per spheroids. The spheroid plate was centrifuged at 50 g for 3 minutes and left in the incubator overnight.

### Spheroid seeding and cell culture maintenance

Just before seeding, the diluted laminin coating solution was replaced by the respective maturation medium. Sensory neuron spheroids were seeded within 48 hours after their preparation. One spheroid was placed in the central well of each of the five circuits in the microstructure using a 10 *µ*L pipette. In this process, it is essential to not touch the microstructure or the upper well, as it may cause its detachment from the substrate. Culture medium was changed a few hours after seeding. For the sensory neurons, after 3 days, the Senso-MM maturation medium was changed to Chrono™ Senso-MMx2 Maturation Media (Anatomic Incorporated, cat. 7012), and after two weeks, to a custom-made medium. The custom-made medium was composed of Neurobasal (NB) (Gibco™) medium, to which 5 % of B27 supplement (17504-044), 1 % GlutaMAX (35050-061) and 1 % Pen-Strep (15070-063, all from ThermoFisher) were added. Growth factor supplements were freshly added to 10 mL of neurobasal medium to obtain the following final concentrations: 50 ng/mL of nerve growth factor (NGF, 450-01), 20 ng/mL of brain-derived neurotrophic factor (BDNF, 450-02), 20 ng/mL of glial-derived neutrophic factor (GDNF, 450-10), 20 ng/mL NT-3 (450-03, all from PeproTech) and 5 *µ*M of forskolin (66575-29-9, Sigma Aldrich). Half of the medium was changed twice a week. Dorsal horn neuron spheroids were seeded within 48 hours after preparation. Before seeding, the medium in the upper well at the “D-HORN” compartment was replaced by 5 *µ*L of Matrigel^®^ (Corning^®^, 354230) diluted 1:1 in cold DHN-MM maturation medium. After incubating the sample for at least 5 minutes at 37 °C to ensure crosslinking of the Matrigel^®^, approximately 10 hiDHN spheroids were seeded in the well at the “D-HORN” compartment. Culture medium was added a few hours after seeding. To not disturb the hiDHN spheroids, medium in the well was not aspirated but 10 *µ*L of DHN-MM medium was added to the well three times per week.

### HPEK seeding and maintenance

Juvenile HPEKs, isolated from pooled donors, were acquired from CELLnTEC (CELLnTEC Advanced Systems AG) at passage 4. HPEKs were plated for expansion in a T25 flask at seeding density 4000 cells/cm^2^, maintained in CnT-07 Epithelial Proliferation Medium (CELLnTEC Advanced Systems AG), then detached with Accutase (CnT-Accutase-100, CELLnTEC) and stored in liquid nitrogen in Bambanker freezing medium (Nippon Genetics, BB02). A cryovial was removed from storage and thawed in a 37 °C dry bath until there was only a small piece of ice left. After taking the vial into a laminar flow hood, the cells were transferred into a centrifuge tube and diluted with twice the amount of warm CnT-07, and subsequently diluted to 160 000 cells/mL. Before seeding, the medium in the upper well at the “SKIN” compartment was replaced by Matrigel^®^, diluted to 1:25 in cold CnT-07. After at least 5 minutes of incubation at 37 °C, the Matrigel^®^ coating was as-pirated and 15 *µ*L of the HPEK suspension was seeded per well, resulting in a seeding density of 80 000 cells/cm^2^. The medium in the well was changed a few hours after seeding. Medium was changed three times per week by replacing 10 *µ*L of CnT-07 in the well.

### HD-MEA chip handling

Electrophysiological recordings were performed in an incubator with 35°C air temperature, 90% humidity and 5% CO_2_ concentration; this setup allowed for continuous recordings up to several hours. After placing the MEA chip on the recording unit, it was allowed to rest for at least five minutes before a recording session was initiated, ensuring that any spontaneous activity disrupted by movement or changes in CO_2_ levels and temperature could return to baseline. Electrophysiology data collection (spontaneous and stimulation-based recordings) was performed within 24 hours after the last medium change to limit variability due to changes in ion concentration and acidity.

### Circuit selection for data collection

Circuits were selected based on the previously mentioned impedance map. A threshold was applied to identify electrodes not covered by the microstructure. A selection of about 1000 electrodes for each circuit from which to record electrical signals was performed using a custom-built previously published software ^**?**^. Selected electrodes were then routed to available amplifiers and the resulting configuration was downloaded to the chip using the Python application programming interface (API) provided by the chip manufacturer (MaxWell Biosystems).

### Recording of electrophysiological voltage traces

The circuit activity was tracked by recording the voltage on the routed electrodes. Voltage recordings were acquired at 20 kHz sampling frequency with a resolution of 10 bits and a recording range of approximately ± 3.2 mV, which results in a least significant bit (LSB) corresponding to 6.3 *µ*V. Using custom software based on the MaxWell Python API, the raw traces were recorded and stored as HDF5-files. Spontaneous activity was recorded for 2 minutes per circuit.

### Voltage- and frequency stimulation

To apply a repetitive stimulation pattern to a circuit, a custom Python script was created. The script makes use of the MaxWell Python API to send commands to the system hub. One stimulation electrode per microchannel was selected based on an activity map during which spontaneous activity over all electrodes was recorded for 1 minute. For each microchannel, the electrode with the highest recorded spike amplitude in the outermost 250 *µ*m of the recording area was selected as the stimulation electrode, implying good signal to noise ratio and the presence of an axon. If no electrode was found in the outermost 250 *µ*m, incremental regions of 250 *µ*m were searched. An example of the electrode selection can be found in Fig. S10 (Supporting Information). Stimulation consisted in 100 pulses applied at the following frequencies: 16 Hz, 32 Hz, 64 Hz, and 96 Hz, in a randomized order. A biphasic pulse with a leading cathodic phase and 400 *µ*s pulse width was used. Each of the 15 microchannel was sequentially stimulated one side after the other at a voltage of 1000 mV. The amplitude was rounded to the closest value available on the 10-bit DAC with approximately 3.2 mV step size.

### Image acquisition and analysis

An inverted confocal laser scanning microscope (FluoView 3000, Olympus) was used to image the fluorescently labeled cultures. Channels typically acquired were 488 nm (Calcein AM 1:2000 in PBS or green CMFDA), 405 nm (Violet Calcein AM, 1:1000 in PBS), 561 nm (Red-Orange Calcein AM 1:1000 in PBS) and phase contrast brightfield images (only for glass bottom dishes). The medium of the central compartment was replaced by the staining solution and incubated for 1 hour and 15 minutes before rinsing with warm PBS. Then, staining solutions were added for 25 minutes incubation in the lateral compartments without rinsing, to avoid disturbing cells. Fluorescence imaging of the CMOS chips was done by removing all excess medium and using a round glass coverslip (10 mm diameter) mounted on top of the PDMS microstructure. The surface tension between the cover slip and the chip enabled us to invert the chip for imaging in an inverted microscope. For mounting in the CLSM, the CMOS chip was placed in the recording unit which in turn was mounted into a custom made metal insert that fits into the stage of the microscope. This configuration enabled inverted mounting for imaging ^37^. Microscope images were processed using Fiji ^**?**^. Importantly, due to their size, stained somata are brighter and thus more visible than axons on microscopy images. To enhance the intensity of the axons compared to the somata, a pixel logarithm operator was applied to all the representative fluorescent images shown in the figures of this paper. The brightness and contrast were manually adjusted to suppress background fluorescence.

### Raw data processing

Raw data were processed using a band-pass filter (4th order acausal Butterworth filter, 300-3500 Hz). The baseline noise of the signal was characterized using the median absolute deviation for each electrode ^**?**^. Action potential event times were extracted from the voltage trace by identifying negative signal peaks below a threshold of 10 times the baseline noise. Successive events within 0.75 ms were discarded to avoid multiple detection of the same spike. Spike amplitudes were defined as the absolute value of the negative amplitude of the detected peak. Spike waveforms were extracted from the filtered voltage trace using the data within a −1.5 ms to 1.5 ms window around the timestamp of the detected spike.

### Individual microchannel identification

Every single microchannel is physically spanned by a set of electrodes, which results in row-like arrangement of pixels on the impedance map representation of the microstructure. Therefore, microchannels were visually identified on the impedance scan and respective electrodes/pixels were manually selected via a custommade GUI ^**?**^ and saved in a separate file. An example of the resulting microchannel identification is shown in Fig. S11 (Supporting Information).

### Individual axon identification with spike sorting

As multiple axons grow within one microchannel, spike sorting was performed to identify individual axons. For each microchannel, all spanning electrodes were ranked based on the their mean detected spike amplitude and firing rate. The electrode with the highest combined rank, where spike amplitude has weight 0.6 and firing rate weight 0.4, was selected as reference electrode. Voltage traces were then extracted around the detected spike timestamps of the reference electrode for all the other electrodes. Two additional reference electrodes were selected based on the same metric, with a constraint on the minimal distance between the three reference electrode of at least 100 *µ*m, to maximize the information regarding spike waveform propagation by avoiding redundancy of neighboring electrodes. Principal component analysis (PCA) was applied to the original feature space consisting of concatenated mean waveforms of the three reference electrodes, and subsequently, clustering was performed on the first three principal components using a Bayesian Gaussian Mixture Model (Python scikit-learn library) with a maximum of 10 components. To avoid over-clustering, spike clusters were then merged. Clusters with high internal variability, defined in Equation (1) were not merged.

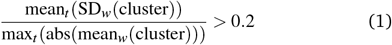

Here *x*_*t*_ represents an operation over the time axis, *x*_*w*_ represents an operation over the waveforms in the cluster, and “cluster” represents all concatenated waveforms of the three reference electrodes. Furthermore, for each cluster, the direction of propagation was determined by detecting the spike on the mean waveform on the reference electrodes. If directionality could not be determined, the cluster was not merged. For all clusters eligible for merging, the cosine similarity matrix was calculated. The merging sequence was determined based on highest cosine similarity. Merging of two clusters was executed if they had the same direction of propagation, the cosine similarity of the concatenated mean waveforms of the three reference electrodes exceeded 0.9 and the distance in the linear discriminant analysis (LDA) space was lower than 5. Each resulting spike cluster was considered to be representative of an axon. Identified axons were discarded if they contained less than 10 spikes, corresponding to a firing rate of 0.0833 Hz.

### Spontaneous activity analysis

To quantify the number of sorted active axons per microchannel, only microchannels with at least one identified axon were considered, to avoid the effect of improperly fabricated or clogged microchannels. Firing rate of the sorted active axons was extracted based on the number of detected spikes at the first reference electrode. For all electrodes in the microchannel, waveforms for each identified axon were extracted around the timestamps of the spikes detected on the first reference electrode. For each electrode, spike detection was applied on the mean waveform. This allowed to visualize the propagation of the waveform along the microchannel. The velocity, and thus the conduction speed and directionality of the waveforms for each identified axon was extracted using a linear fit on the position and spike time over all electrodes in the microchannel with a detectable spike. Only fits with a Pearson correlation coefficient above 0.9 were retained. Axons were also discarded if less than 5 electrodes had a detectable spike in their mean waveform. Metrics extraction is shown in Fig. 5C. Data consisted in 2 × 60 microchannels distributed across four circuits on one chip for the hiSN-only cultures, 2 × 75 microchannels distributed across 5 circuits on one chip (co-culture of hiSNs and HPEKs) and 2 × 45 microchannels distributed across 3 circuits on one chip (co-culture of hiSNs and hiDHNs).

### Stimulation-induced response analysis

Stimulation-induced responses were analyzed at two locations: first, at the control electrode, in the stimulated microchannel close to and upstream of the central compartment, second, at the target electrodes, in the microchannels downstream of the central compartment, on the side of the microstructure that was not stimulated. The control and target electrodes were selected based on the previously defined activity map. For each microchannel, the control electrode was defined as the electrode with the highest recorded spike amplitude in the 250 *µ*m closest to the central compartment. If no electrode was found in this region, incremental regions of 250 *µ*m were searched. The target electrode was defined as the electrode with the highest recorded spike amplitude in the complete microchannel. Per stimulation, *i*.*e*., per stimulated microchannel, a post-stimulus raster plot was generated with spikes detected in the control and target electrodes to visually inspect the response of axons in each microchannel to the stimulation, as described previously ^39^. As the stimulation signal saturates the amplifiers capped at 3.2 mV independently from the stimulation amplitude, an artifact amplitude threshold of 2.5 mV was established to detect the stimulation pulse times on the stimulation electrodes. Then, for the control electrode of the stimulated microchannel and the target electrode of each microchannel downstream of the central compartment, spikes were detected in a time window from 0.5 ms to 8 ms after the stimulation artifact. This time window excluded the stimulation artifact and was sufficiently long for the action potentials to travel from the stimulation electrode to the microchannels on the other side of the platform. In order to identify the stimulation-induced responses, the post-stimulus raster plot was analyzed, as previously described and shown ^39^. The raster plot, containing the time of each recorded spike, is a two-dimensional array image in which the spontaneous random spikes correspond to background noise and the stimulation induced spikes form “bands” which are the features of interest. In order to remove the background noise, a bilateral filter was applied (Python OpenCV library). A clustering algorithm (hierarchical clustering, Python scikit-learn library) was then applied to separate and identify the “bands”. It is important to note that the metrics extraction is affected by the band detection method, and that specific corner cases arise and have an impact on the variability. Each “band” was identified as a stimulation-induced response.

Stimulation cannot be applied sufficiently localized to specifically target one microchannel, possibly also stimulating neighboring microchannels, as illustrated in Fig. S10 (Supporting Information). To account for this limitation, responses to the sequential stimulation of each single microchannel were aggregated per stimulation side in the model. A microchannel was considered to be activated when at least one stimulation-induced response was detected for its selected electrode. At the control, only responses to stimulation in the same microchannel were considered. Per target microchannel, responses to stimulations in all of the microchannels on the opposite side were aggregated, and a microchannel was considered activated when at least one response was detected across all stimulations. The number of activated microchannels was normalized by the total number of microchannels per side and expressed as a percentage. There was not error bar for standard error or mean for this metric. Other metrics were extracted per stimulation-induced response and included the point of conduction failure, defined as the the ratio of the last stimulation pulse to elicit a response and the total number of stimulation pulses, and the response probability, defined as the ratio of the amount of detected spikes in the response and the total number of stimulation pulses. The response probability was further normalized to the difference between the first and final stimulation pulse that elicited a detectable response, to eliminate the effect of conduction failure. Metrics extraction was previously shown ^39^. Data for the stimulation-induced response analysis was collected for the same samples as for the spontaneous activity analysis.

### Statistical Analysis

Data was considered non-normally distributed based on the Shapiro-Wilk test for normality with a significance level of *p* = 0.05. For comparison of two samples, the Mann-Whitney U test was applied to a significance level of *p* = 0.05. Metrics are reported based on the mean and the standard error of mean where possible. All statistical tests were performed using the Python SciPy Stats library.

## Acknowledgments

This research was supported by ETH Zürich, the Swiss National Science Foundation (SNSF) (project number 182779) and the Human Frontier Science Program (HFSP) postdoctoral fellowship.

## Author contributions

BFC conceptualized and designed the research project. BFC, TO and CZ wrote the initial draft of the manuscript. BFC performed preliminary experiments. BFC and CZ designed the microfluidic device, with inputs from TR. BFC, TO, CZ conducted the experiments, collected the data and analyzed the data, with support from BFC. GP helped with HPEK experimental methods and result interpretation. JV secured funding for the projects. All co-authors reviewed and approved the manuscript.

## Supporting information

Supporting information associated with this article can be found in the online version here

