## Supplementary Information for "A three-compartment microfluidic platform for investigating signal transmission in the human sensory pathway^†^"

### S1 Supporting Information

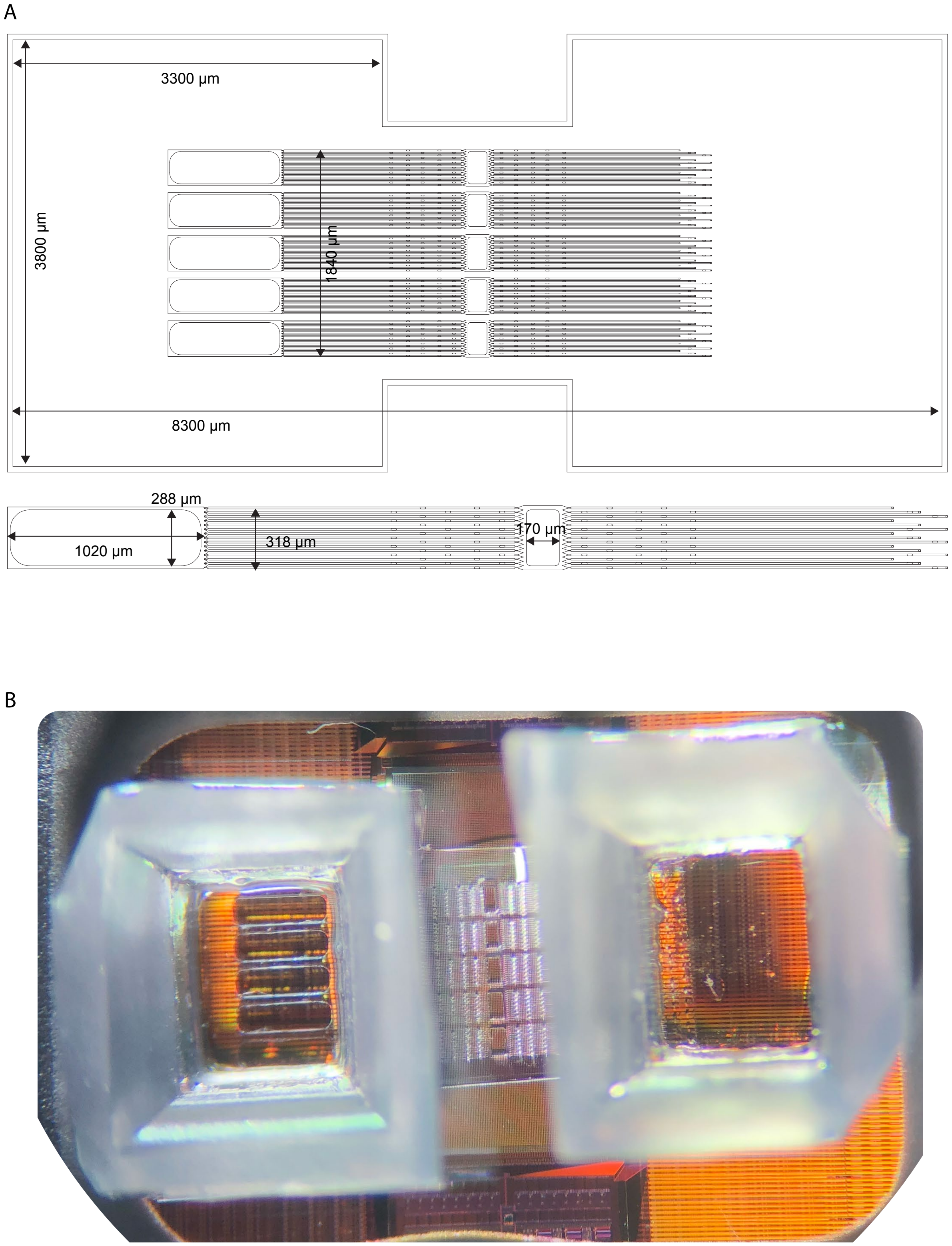


**Figure S1:** A: Dimension details of the three-compartment PDMS microstructure design. The original AutoCAD file can be obtained upon request. B: Picture of the platform on top of the HD-MEA taken through a stereo-microscope, in air.

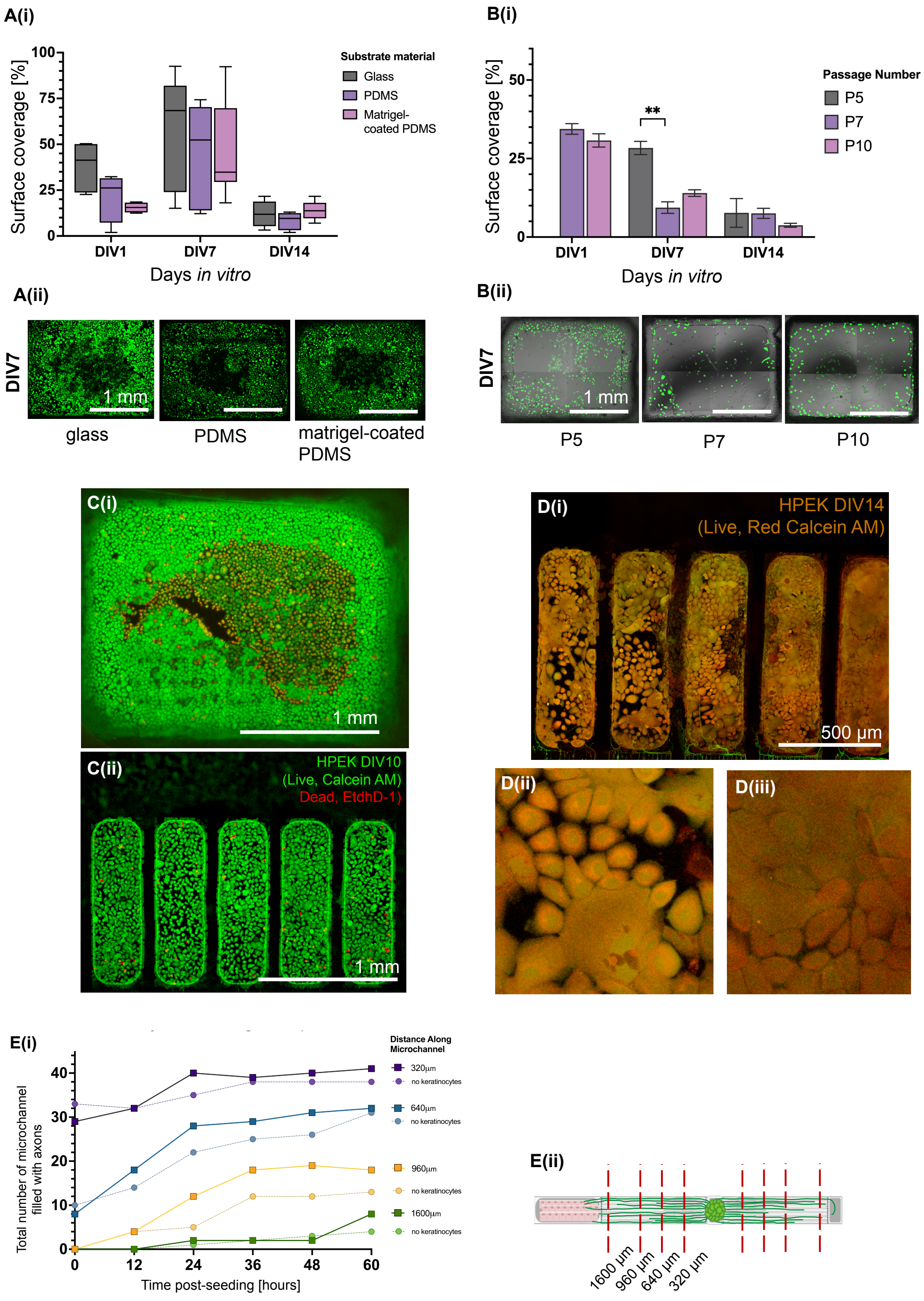


**Figure S2: Effect of HPEK culture conditions.** **A**(i): Quantified surface coverage of HPEK seeded at a different passage numbers. N=4 samples for each bar and $p < 0.0001$ at DIV 7 (Kruskal-Wallis test). (ii): Representative images for each conditions taken at DIV 7. **B**: Quantified surface coverage of HPEK seeded on different substrate materials. N=4 to 9 for each box. For A and B, unless shown to be an independent variable, default conditions were a seeding density of 80,000 cells/cm$^2$, seeded at passage 6 and on uncoated glass. (ii): Representative images for each conditions taken at DIV 7. **C**: Live-dead staining for HPEK culture at DIV 10 in single PDMS well compartment (i) and in five smaller sub-compartments (ii). Cells start dying and detach from the center of the culture area, which is improved in the sub-compartment design. **D**: HPEK morphology is variable across sub-compartments in the same PDMS well, ranging from a proliferative state (ii) to a seemingly monolayer formation phenotype (iii). **E**(i): Comparison of axon growth rate towards compartments with and without HPEKs across different time points. Data is shown for number of "filled"[2]microchannels with axons at given distance cross-points as illustrated in E(ii) for a total of 45 microchannels distributed across 3 circuits.

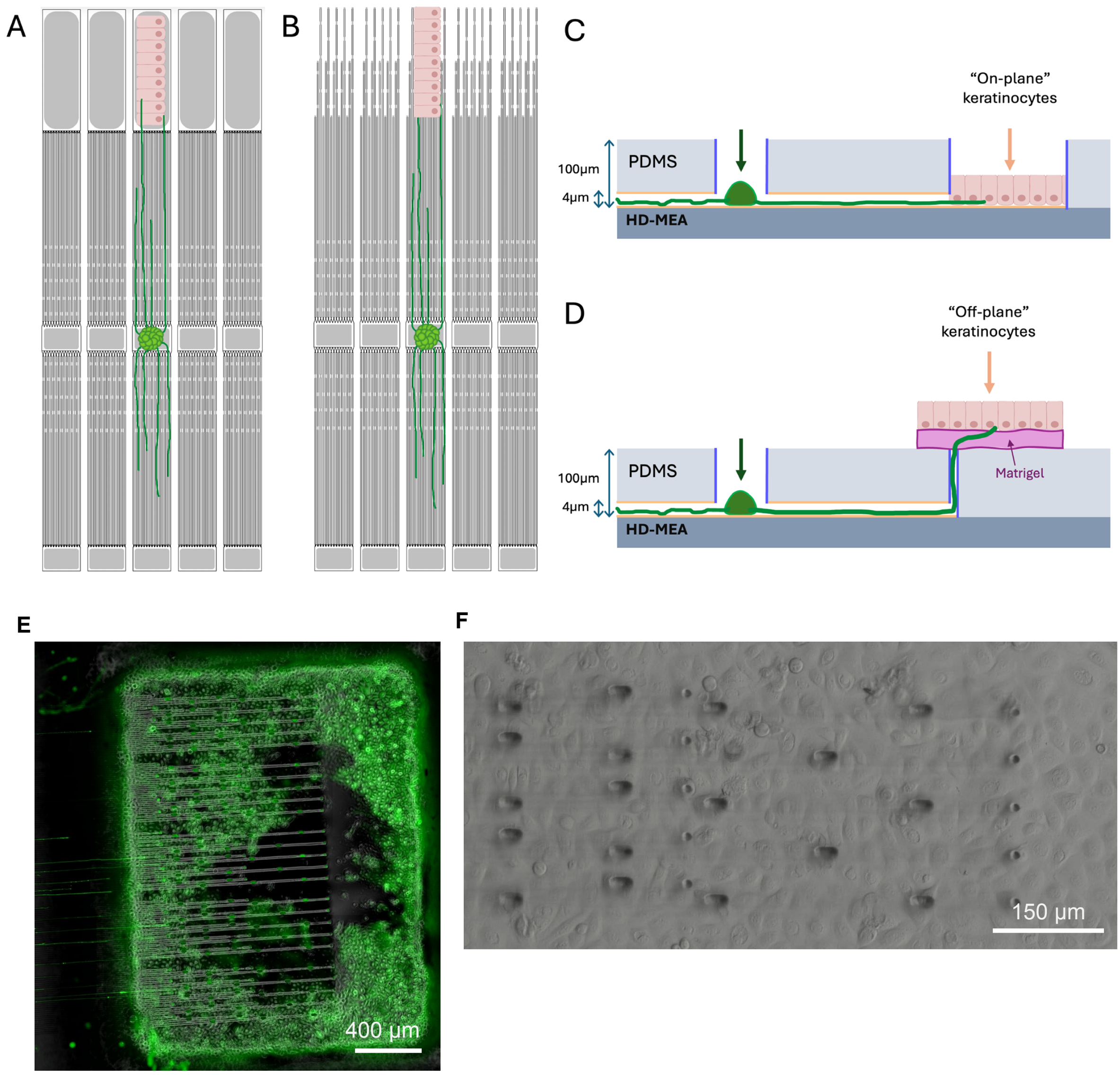


**Figure S3: Illustration of two strategies ("in-plane" and "off-plane") for the neurons to interface HPEKs.** **A** and **C**: Top (A) and cross section (C) view schematic of the in-plane approach. **B** and **D**: Top (B) and cross section (D) view of the "off-plane" design. In the "off-plane" design, HPEKs are seeded on top of the PDMS microstructure, to encourage axon growth in an upwards direction. A Matrigel coating can be employed to facilitate axon traversal to access more HPEK cells. **E**: Fluorescence image showing feasibility of HPEK culture on the top plane of the PDMS microstructure. The focus is at the plane of the microchannels where visible axons grew until the upward wells. Live cells were stained with Calcein AM. **F**: Bright-field image of the top plane with HPEKs.

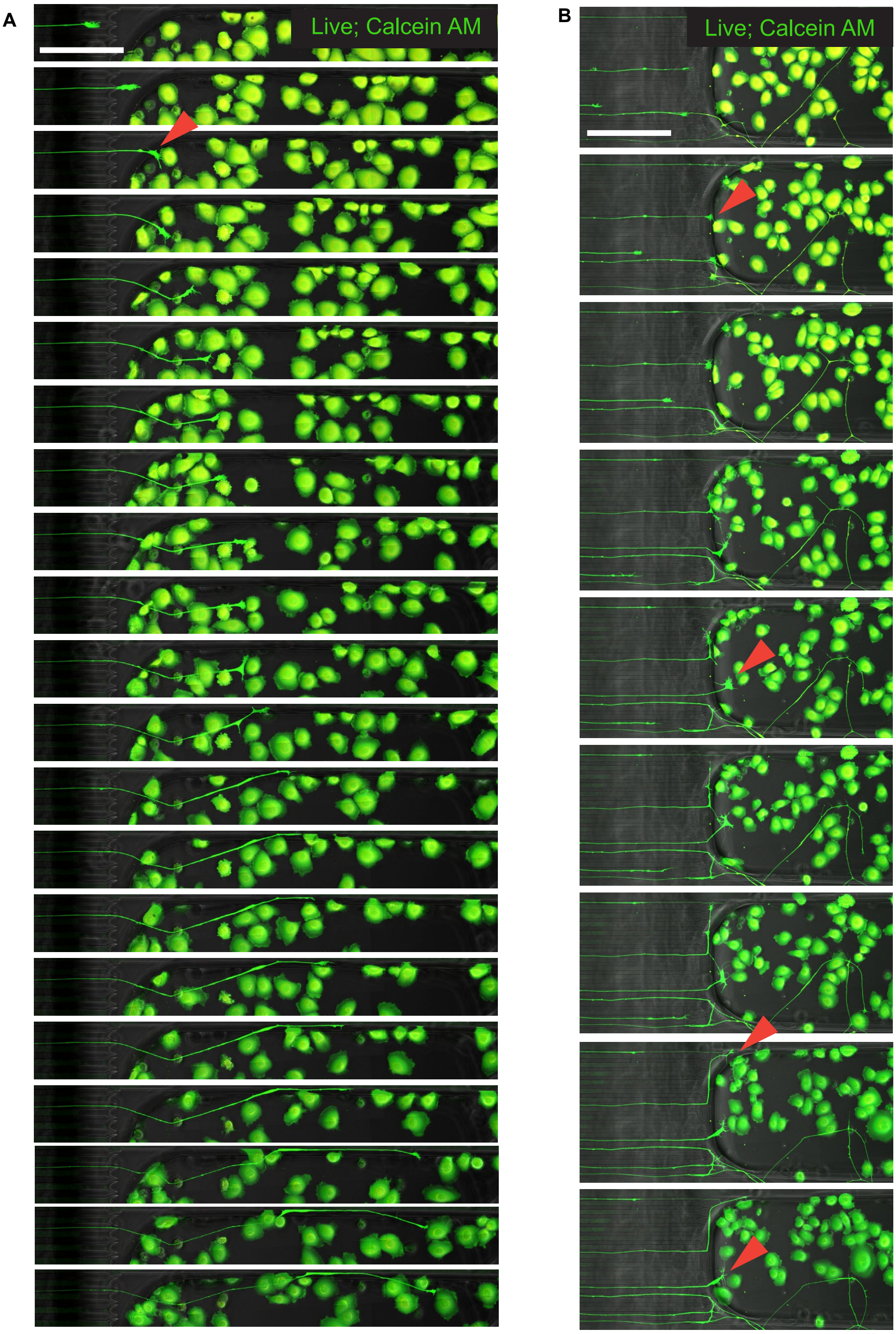


**Figure S4: Time-lapse of axons and HPEKs at early seeding stage.** **A** and **B**: Time-lapse images showing one axon (A) or more axons (B) at DIV 4 growing inside the "SKIN" compartment and interacting with HPEK cells (DIV 1). Elapsed time between each graph is 30 minutes. Both cell types were stained with Calcein AM. Scale bars = 100 µm in A and 150 µm in B.

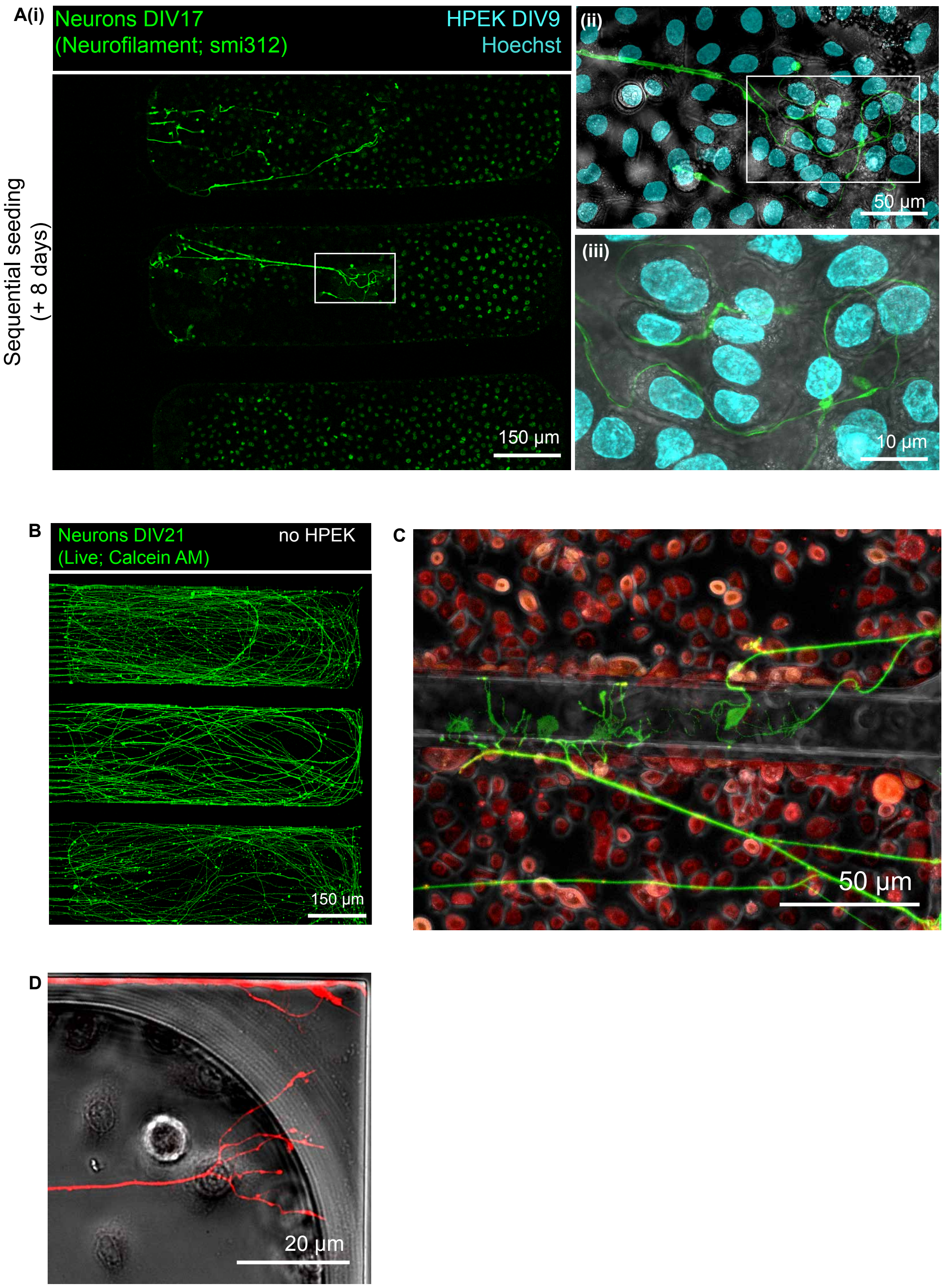


**Figure S5: hiSN axons and HPEKs interface in the "SKIN" compartment A:** Fluorescence image of the keratinocyte compartment with incoming axons from microchannels when HPEKs were seeded 9 days after neuron seeding. Cells were fixed and stained for neurofilament (smi312, green) and Hoechst (cyan). **B:** "SKIN" compartment without any HPEKs seeded and hiSN axons grown at DIV 21. **C:** Zoomed-in view from main figure 2E(ii) showing hiSN growing between HPEKs. Some hiSN axonal ends grow below PDMS inter-subcompartment walls and some exhibit a nerve-ending-like morphology. **D**: Another example of nerve-ending like morphology.

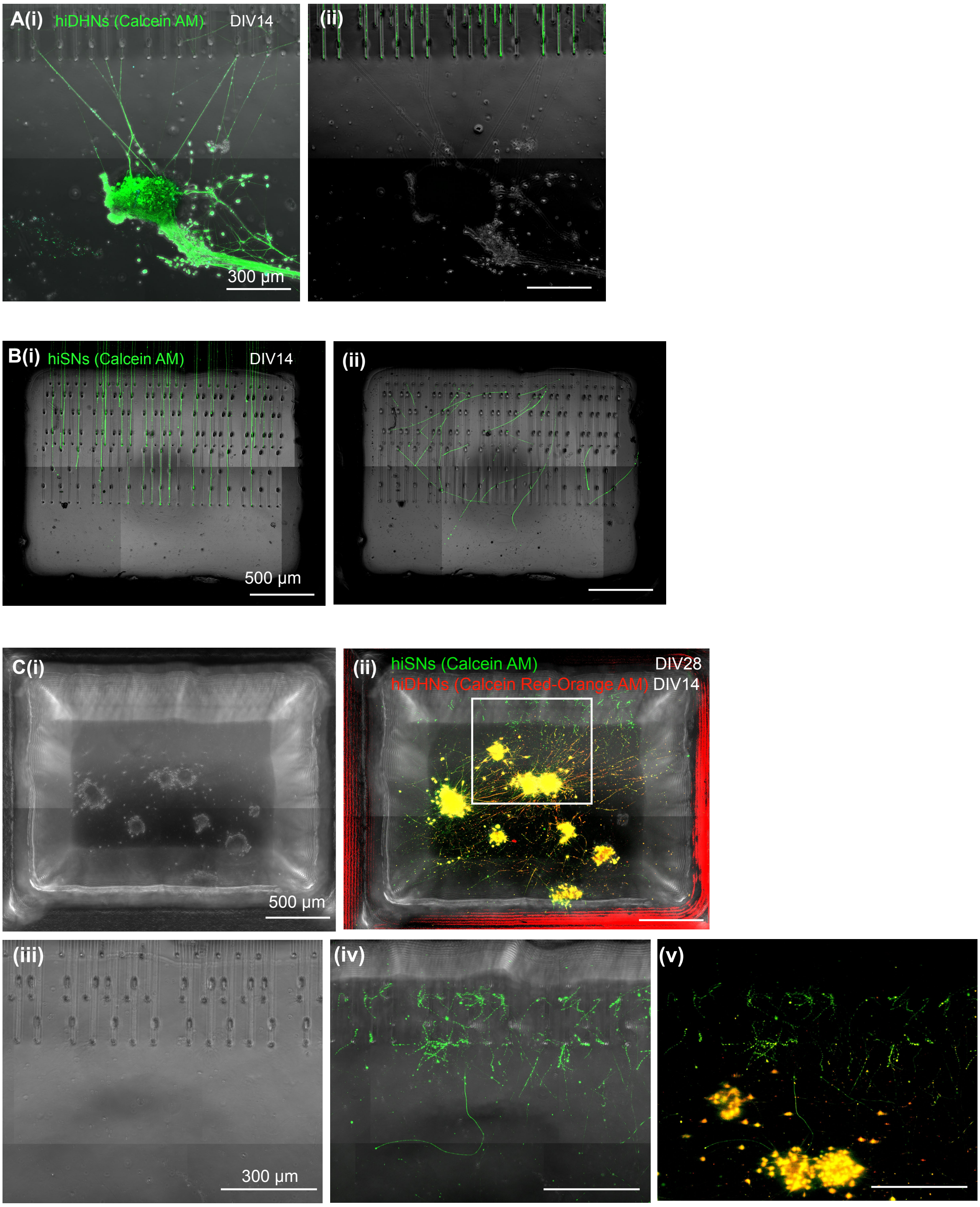


**Figure S6: Optimization of hiDHN culture in "D-HORN" compartment. A:** Fluorescence image of (i) top of PDMS microstructure ("off-plane") where hiDHN spheroids were seeded (without Matrigel) and (ii) microchannel plane where some hiDHN axons grew in, showing that when seeded hiDHNs directly on the "off-plane", axons extend to find irregularities from the upward microwells and grow in, that would interfere with sensory neuron growth. Taken at DIV 14. **B:** Fluorescence image of (i) microchannel plane where some hiSNs were seeded in the center (not shown) and (ii) top of PDMS microstructure ("off-plane") where hiSN axons grew upward, showing the feasibility of the "off-plane" approach to axons from the bottom plane growing up in the compartment, that is leveraged in the platform. Image taken at DIV 14. **C:** Maximum projection of z-stack microscopy of the DH compartment from "off-plane" level (top of microstructure) to top of Matrigel (about 400 µm higher up. (i) bright-field and (ii) composite image of the DH compartment showing hiDHN spheroids seeded higher up on Matrigel. (iii-v) are zoomed-in views of (ii) showing the hiSN axons (green) and hiDHN spheroids (red-orange) interfacing in the DH compartment. hiSNs: DIV 28; hiDHNs: DIV 14.

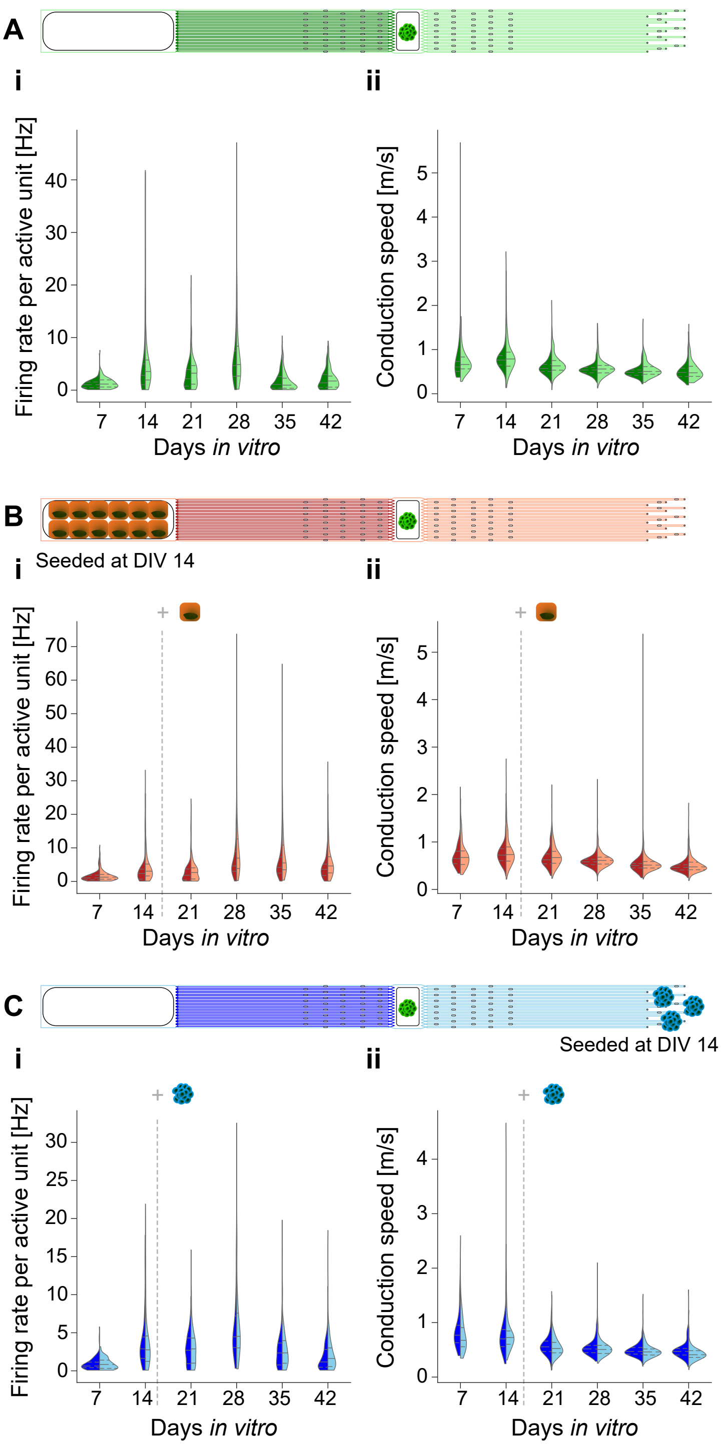


**Figure S7: Distribution of spontaneous parameters per model side across DIVs A:** Distributions for a hiSN-only culture ($2 \times 60$ microchannels distributed across 4 circuits on one chip). i: Firing rate per active unit, smoothed distribution over active units is plotted. ii: Conduction speed, smoothed distribution over active units is plotted. **B:** Distributions for a co-culture of hiSNs and HPEKs ($2 \times 75$ microchannels distributed across 5 circuits on one chip). i-ii: Panels are identical to A: i-ii. **C:** Distributions for a co-culture of hiSNs and hiDHNs ($2 \times 45$ microchannels distributed across 3 circuits on one chip). i-ii: Panels are identical to A: i-ii. In co-culture conditions, HPEKs and DH neurons were seeded at DIV 14.

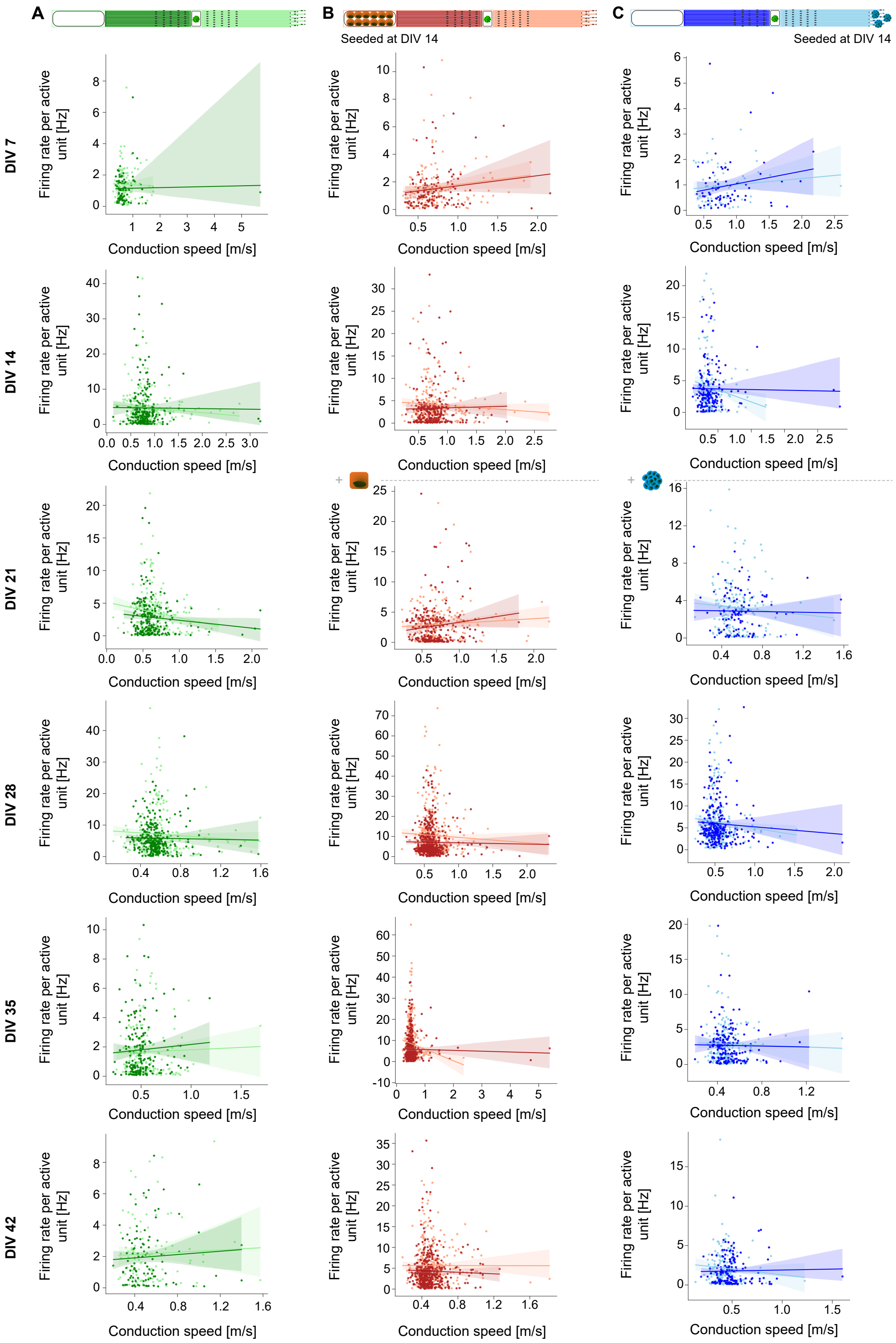


**Figure S8: Correlation of spontaneous firing rate and conduction speed per active unit over co-culture conditions and DIV. A:** hiSN-only culture (2 × 60 microchannels distributed across 4 circuits on one chip). **B:** Co-culture of hiSNs and HPEKs (2 × 75 microchannels distributed across 5 circuits on one chip). **C:** Co-culture of hiSNs and hiDHNs (2 × 45 microchannels distributed across 3 circuits on one chip). Linear fit and 95%-confidence interval are shown. In co-culture conditions, HPEKs and DH neurons were seeded at DIV 14.

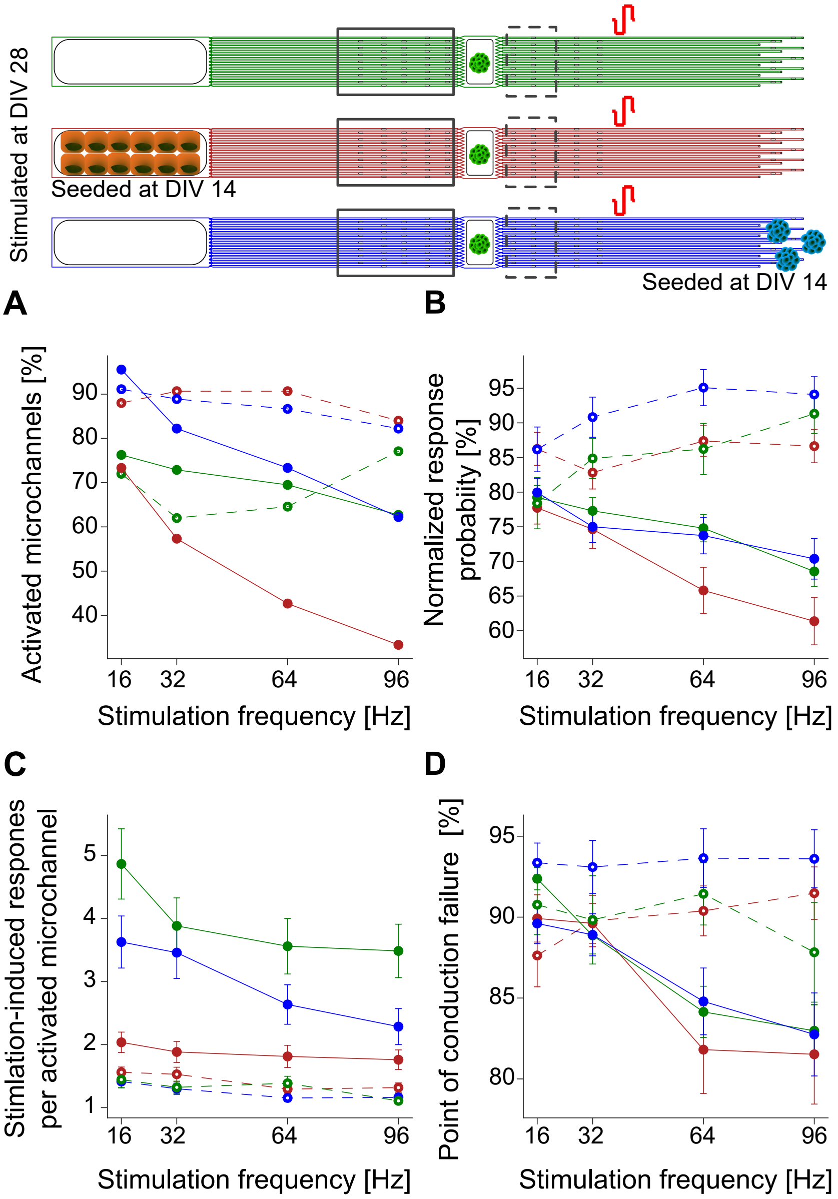


**Figure S9: Frequency-dependency of transmission of stimulation-induced responses for stimulation at "D-HORN" side.** Quantification of response parameters for sequential stimulation with different pulse frequency at the DH side at DIV 28 for the following conditions: 1) hiSN-only culture (2 × 60 microchannels distributed across 4 circuits on one chip), 2) co-culture of hiSNs and HPEKs (2 × 75 microchannels distributed across 5 circuits on one chip) and 3) co-culture of hiSNs and hiDHNs (2 × 45 microchannels distributed across 3 circuits on one chip). In co-culture conditions, HPEKs and hiDHNs were seeded at DIV 14. **A:** Percentage of microchannels that are activated, *i.e.*, have at least one detected response. **B:** Normalized response probability as percentage of total applied stimulation pulses, means and standard error of means over responses are plotted. **C:** Number of detected stimulation-induced responses per activated microchannel, means and standard error of means over activated microchannels are plotted. **D:** Point of conduction failure as percentage of total applied stimulation pulses, means and standard error of means over responses are plotted.

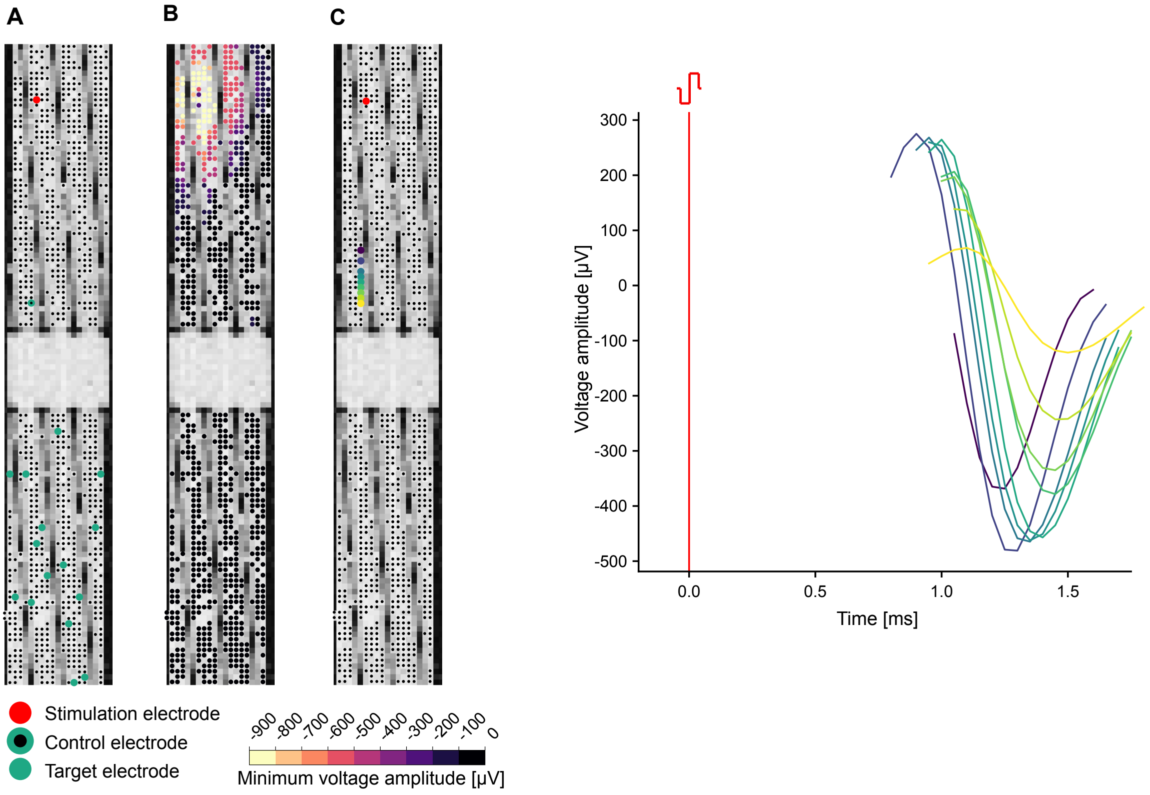


**Figure S10: Voltage stimulation. A:** Electrode selection for voltage stimulation and subsequent analysis. **B:** Minimum detected voltage amplitude during stimulation, representative of the spread of the stimulation artifact, and thus indicating the location of stimulation. **C:** Waveforms recorded along a microchannel after successful stimulation.

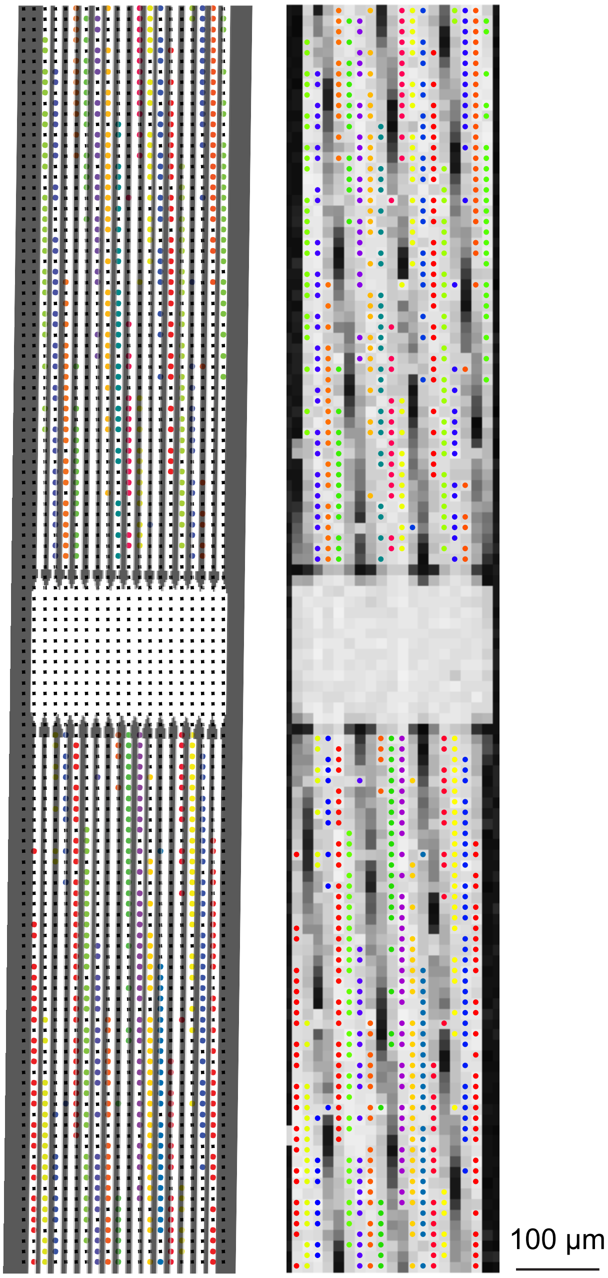


**Figure S11: Microchannel identification.** Identification of microchannels on recording electrodes is performed based on a voltage map. Each group of colored electrodes represents a microchannel.
